# SOX2 drives fetal reprogramming and reversible dormancy in colorectal cancer

**DOI:** 10.1101/2024.09.11.612412

**Authors:** Anna Baulies, Veronica Moncho-Amor, Diana Drago-Garcia, Ania Kucharska, Probir Chakravarty, Manuel Moreno-Valladares, Sara Cruces-Salguero, Florian Hubl, Colin Hutton, Hanue Kim, Ander Matheu, Robin Lovell-Badge, Vivian S. W. Li

## Abstract

Cellular plasticity plays critical roles in tissue regeneration, tumour progression and therapeutic resistance. However, the mechanism underlying this cell state transition remains elusive. Here, we show that the transcription factor SOX2 induces fetal reprogramming and reversible dormancy in colorectal cancer (CRC). SOX2 expression correlates with fetal reprogramming and poor prognosis in human primary and metastatic colorectal adenocarcinomas. In mouse CRC models, rare slow-cycling SOX2^+^SCA1^+^ cells are detected in early *Apc*-deleted mouse tumours that undergo slow clonal expansion over time. In contrast, the SOX2^+^ clones were found proliferative in advanced *Apc^-/-^;Kras^G12D/+^;p53^-/-^;Tgfbr2^-/-^*(AKPT) tumours, accompanied by dynamic cell state reprogramming from LGR5^+^ to LGR5^-^SCA1^+^ cells. Using transgenic mouse models, we demonstrate that ectopic expression of SOX2 inhibits intestinal lineage differentiation and induces fetal reprogramming. SOX2+ cells adopt dynamic cell cycle states depending on its expression level. High SOX2 expression results in hyperproliferation, whereas low SOX2 levels induce senescence-mediated dormancy. We find that loss of p53 can reverse SOX2-induced senescence, in line with the dormant cell state exit of the SOX2+ cells observed in advanced tumours. Finally, SOX2 expression is induced by 5-FU treatment in CRC. SOX2-expressing organoids exhibit increased tolerance to chemotherapy treatment, whilst deletion of SOX2 in AKPT tumour organoids sensitises drug responses. We propose that SOX2-induced plasticity and reversible dormancy promotes tumour progression and drug tolerance in CRC.

## Main Text

Cellular plasticity refers to the capability of cells to be reprogrammed and to acquire a new cell state, which can be triggered by genetic/epigenetic-dependent intrinsic mechanisms or in response to external stimuli without genetic mutations^1^. In the intestine, cellular plasticity has been demonstrated in injury-induced regeneration and in cancer. Several studies have previously reported that the intestinal epithelium undergoes reprogramming upon infection, irradiation or colitis^2–5^. This allows transient reversion of committed adult intestinal cells back to regenerative or fetal-like cell states to facilitate regeneration. Similarly, phenotypic plasticity has also been reported in cancer and is believed to drive malignant progression and acquire resistance to therapies^6,7^. In colorectal cancer (CRC), LGR5+ cells serve as cancer stem cells with high clonal expansion capacity^8,9^, yet selective ablation of LGR5+ cells induces tumour plasticity to regenerate cancer stem cells from LGR5- cells^9,10^. In addition, disseminated LGR5- CRC cells have also been demonstrated to drive distant metastasis by reacquiring LGR5+ properties at secondary site^11^, indicating that phenotypic plasticity plays a crucial role in metastasis and tumour recurrence after treatment. Interestingly, several studies have recently reported the observation of cellular reprogramming during CRC initiation, progression and drug resistance^12–14^, suggesting a link between tumour plasticity and transcriptional reprogramming. Deciphering the molecular mechanism underlying cellular reprogramming is key to improving cancer patient survival.

The transcription factor sex-determining region Y-box 2 (SOX2) is one of the key pluripotency drivers that plays an important role during early vertebrate embryogenesis and is required later in the developmental process of various organs including the brain, neural tube, germ cells, and the foregut^15–20^. In particular, SOX2 is one of the four Yamanaka factors necessary for reprogramming somatic cells to induced pluripotent stem cells (iPSCs)^21^, and has been shown to regulate pluripotency by transcriptional epigenetic remodelling^22,23^. During embryogenesis, SOX2 plays a critical role in gastrointestinal (GI) tract development and specification. Mice with hypomorphic SOX2 alleles showed foregut defects and altered expression of GI cell type markers^24^. In adult tissues, SOX2 expression is restricted to the stomach for gastric epithelial specification and homeostasis, but it is completely absent in the intestinal epithelium^24^. Whilst SOX2 is not required for adult intestinal homeostasis, aberrant expression of SOX2 has been reported in CRC and was found to be correlated to high tumour grade, metastasis and poor response to neoadjuvant therapy^25–28^, suggesting a role in tumour plasticity and cancer progression. Here, we explore the putative role of SOX2 in tumour plasticity and reprogramming in CRC.

## Results

### SOX2 expression correlates with poor prognosis and reprogramming in human CRC

Aberrant SOX2 expression has been reported in many cancer types and is often associated with poor prognosis^25^. Comparison of the transcriptomic data from a cohort of 607 TCGA colorectal adenocarcinoma (COAD) and rectum adenocarcinoma patient samples with 51 normal colorectal tissues showed that *SOX2* expression is indeed upregulated in CRC, specifically in a sub-group of high *SOX2*-expressing tumoral samples (Fig. 1a). Of note, SOX2 is expressed in the enteric nervous system^29^, hence it is detected in normal colonic tissues. Kaplan–Meier survival analysis demonstrated that high *SOX2* expression correlates with poor survival (hazard ratio (HR) log-rank p = 0.01639, P = 0.014) (Fig. 1b). A recent study has redefined CRC subtypes based on gene ontology and biological activation state: pathway-derived subtype 1 (PDS1) tumours are in a canonical LGR5+ hyperproliferating state with good prognosis; PDS2 tumours are enriched for regenerative states with high stromal and immune signatures; and PDS3 tumours are in slow-cycling states with the worst prognosis^30^. Interestingly, both *SOX2* and its downstream target *SOX21*^31^ are enriched in PDS3 tumours (Fig. 1c and S1a), consolidating the link between high SOX2 expression and poor prognosis. To validate the results, immunostaining of SOX2 was performed in COAD, metastatic primary COAD and normal colonic tissue. As expected, SOX2 was not expressed in normal colonic epithelium (Fig. 1d). On the other hand, expression of SOX2 was readily detected in human COADs, and was highly expressed in metastatic COADs (Fig. 1d). Both SOX2+KI67- and SOX2+KI67+ cells were detected in COAD samples, whilst most of the SOX2+ cells were KI67+ in metastatic primary COADs (Fig. 1d), suggesting that SOX2+ cells proliferate and expand during tumour progression.

**Figure 1.**
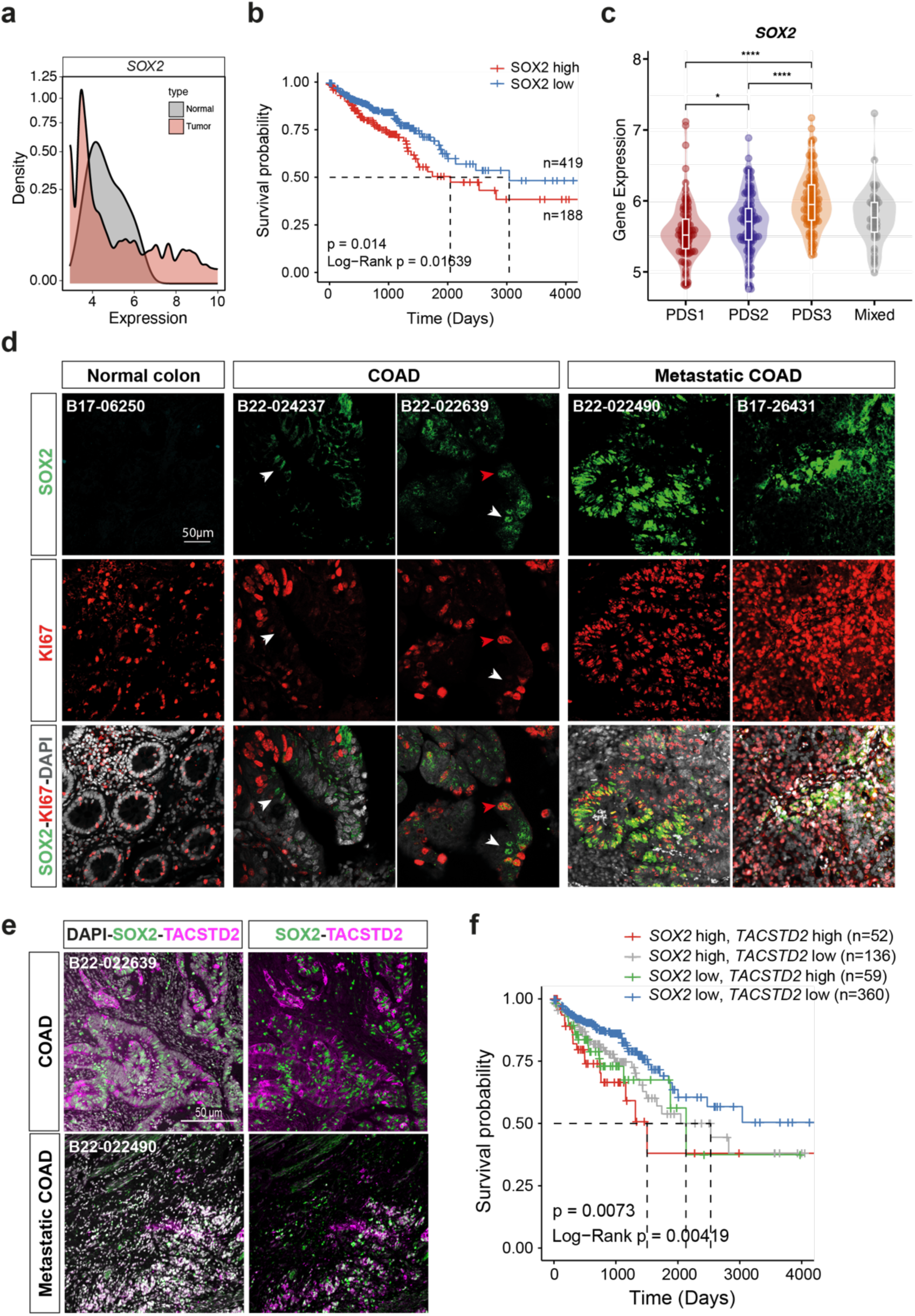
SOX2 correlates with poor prognosis and reprogramming in human CRC. **a**, Density plot of *SOX2* expression in normal, tumour COAD and READ patients from TCGA. **b**, Survival probability for COAD and READ patients from TCGA according to *SOX2* expression status (*SOX2* high n=419, *SOX2* low n=188. Global Log-Rank P-values for the Kaplan-Meier estimates and Cox proportional-hazards model are shown. The hazard ratio is 1.6 with a p-value of 0.015 for high Sox2 expression. **c**, Violin plot displaying *SOX2* expression across PDS in the FOCUS cohort (GSE156915; n=360). **d**, Double immunofluorescence for SOX2 and KI67 in normal colon (normal counterpart of a well differentiated COAD), COAD and Metastatic COAD from patients. Red arrowhead indicates SOX2+KI67_cells; white arrowhead indicates SOX2+KI67- cells. Patient ID indicated at the top. Scale bar, 50μm. **e**, Double immunofluorescence for SOX2 and TACSTD2 in COAD and Metastatic COAD from patients. Scale bar, 50μm. **d,** Double immunofluorescence for SOX2 and SCA1 in APKT lines treated with 5-FU chemotherapeutic drug at the indicated concentrations. Scale bar, 20μm. **f,** Survival probability for COAD and READ patients from TCGA according to *SOX2* and *TACSTD2* expression status in COAD and READ patients from TCGA. (*SOX2* high-*TACSTD2* high n=52, *SOX2* high-*TACSTD2* low n=136, *SOX2* low-*TACSTD2* high n=59, *SOX2* low-*TACSTD2* low n=360). Global Log-Rank P-values for the Kaplan-Meier estimates and Cox proportional-hazards model are shown. Hazard ratios are 1.5 with a p-value of 0.032 for high Sox2 expression; and 1.6 with a p-value of 0.018 for high TACSTD2 expression.

Since SOX2 drives pluripotency, we asked if SOX2 co-localises with reprogramming markers in human COAD. First, we analysed the recently published single-cell and spatial transcriptome analysis of human CRCs^32^ and examined the expression of SOX2, the fetal reprogramming marker Tumor-associated calcium signal transducer 2 (*TACSTD2/TROP2*)^5,33^ and the revival/regenerative marker Clusterin (*CLU*)^2,34^. We found that *SOX2* co-localised with *TACSTD2* in the tumour regions of the CRC tissues, whereas *CLU* localised largely in the paratumour regions (Fig. S1b). The stromal expression pattern of *CLU* suggests that it may not be an appropriate reprogramming marker to represent plasticity in tumour cells. Interestingly, expression of *TACSTD2* is upregulated in human COADs (Fig. S1c) and shows a mild but significant positive correlation with *SOX2* (Fig. S1d). This was confirmed by co-localisation of SOX2 and TACSTD2 proteins in COADs and metastatic COAD tissues (Fig. 1e). Kaplan– Meier survival analysis further shows that *SOX2* high and *TACSTD2* high patients have the shortest median survival (HR log-rank p = 0.00419, P = 0.0073) (Fig. 1f). Together the results indicate that SOX2 expression associates with fetal reprogramming and poor prognosis in human CRC.

### Clonal expansion and fetal reprogramming of SOX2+ cells in mouse adenomas and CRC

To better understand SOX2 expression dynamics during CRC development, we evaluated its expression in mouse premalignant and advanced intestinal tumours. First, we examined the expression of SOX2 in early intestinal tumours using *Lgr5-EGFP-IRES-CreERT2*, *Apc^fl/fl^* mice. Administration of a single low dose of tamoxifen induced adenoma transformation progressively in the intestine. Interestingly, we were able to detect rare SOX2+ cells in small clusters (∼1-3 cells) as early as 23 days post induction (dpi), and the number of SOX2+ cells gradually increased over time (Fig 2a). Notably, at later time points the SOX2+ cells were always found in cluster patterns, suggesting a slow clonal expansion of the SOX2+ cells within the tumours. Importantly, we found that these SOX2+ cells in the early adenomas were mostly non-proliferative (KI67- and EdU-) and expressed the fetal reprogramming marker LY6 member stem cell antigen-1 (SCA1/LY6A)^5,33^ (Fig. 2b-c). This is consistent with the earlier observation that SOX2 is enriched in the slow-cycling PDS3 tumours. Of note, SCA1 expression was detectable in single SOX2+ cells (Fig. 2c), suggesting that SOX2 drives fetal reprogramming cell autonomously.

**Figure 2.**
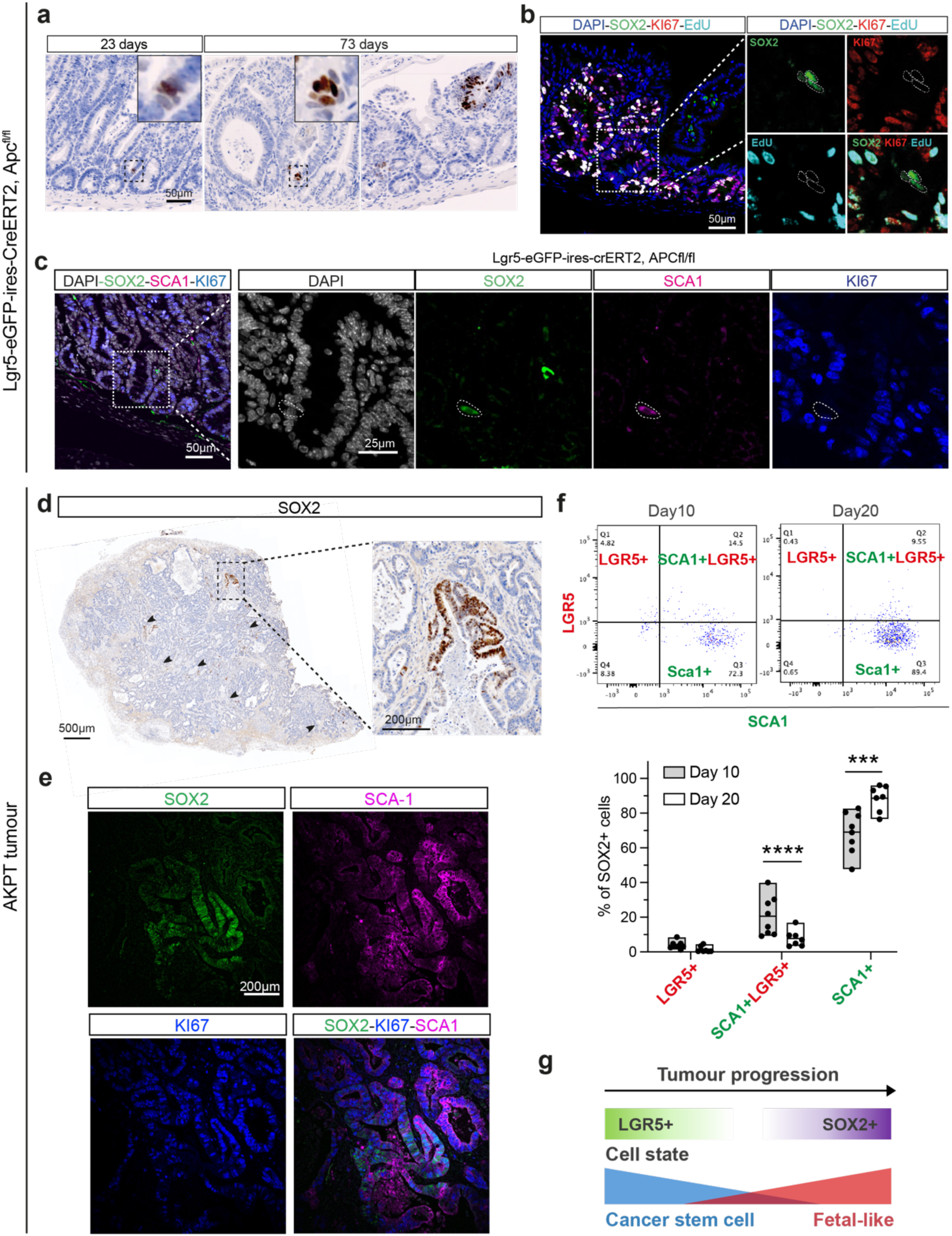
Clonal expansion and fetal reprogramming of SOX2+ cells in mouse adenomas and CRC. **a,** IHC using an antibody against SOX2 on intestinal paraffin sections of Lgr5-eGFP-ires-CreERT2, Apc^fl/fl^ animals at 23 and 73 days after TM treatment. Scale bar, 50μm. **b,** Triple immunofluorescence for SOX2, KI67 and EdU in Lgr5-eGFP-ires-CreERT2, Apc^fl/fl^ animals at 73 days after TM treatment. Scale bar, 50 μm. **c,** Triple immunofluorescence for SOX2, SCA1 and KI67 in Lgr5-eGFP-ires-CreERT2, Apc^fl/fl^ animals at 73 days after tamoxifen treatment. Scale bar, 50μm. **d,** IHC using an antibody against SOX2 on paraffin sections of AKPT tumours. **e,** Triple immunofluorescence for SOX2, SCA1 and KI67 in AKPT tumours. Scale bar, 200μm. **f,** Representative flowcytometry data showing the percentage of Sox2+ cells that are Lgr5+, Sca1+ or both in APKT tumours collected at day 10 (n=8) and 20 (n=7). Data represent mean ± s.e.m. ***P<0.001, ****P<0.0001, 2-way ANOVA. **g**, Schematic illustration summarising the transition of LGR5+ cancer stem cell state to SOX2+ fetal-like cell state during tumour progression.

A recent study reported an aberrant stem cell-like (AbSC) state transition through regeneration and developmental reprogramming in premalignant intestinal tumours using scRNA-seq^35^. To validate our findings, we re-analysed the ATAC-seq and scRNA-seq dataset derived from wildtype (WT) and *Lgr5-Ires-CreRT2*, *Apc^fl/fl^*, *R26^tdT^* (Apc KO) intestine at 28 and 72 dpi to explore *Sox2* expression (Fig. S2a). ATAC-seq analysis on the AbSC population showed increased chromatin accessibility of the *Sox2* gene locus in the Apc KO intestine, suggesting the expression of *Sox2* in this plastic cell population (Fig. S2b). Interestingly, three *Sox2*- expressing cells were detected specifically in this AbSC population at 72 dpi (the total number of cells of this experiment is 916), suggesting a link between SOX2 and reprogramming in early premalignant transformation (Fig. S2c). The absence of *Sox2* expression at the earlier time point (28 days) is likely due to the limitation of the scRNA-seq for detecting rare cell populations. Expression analysis of these three SOX2+ cells revealed absence of the intestinal stem cell (ISC) marker *Lgr5* and *Olfm4* and upregulation of its downstream target *Sox21* as well as the reprogramming markers *Ly6a* (*Sca1*) and *Clu* (Fig. S2d). The results suggest that early adenoma transformation driven by APC loss induces SOX2 expression and cellular reprogramming in a rare slow-cycling cell population.

Next, we characterised SOX2 expression in an advanced CRC model via *in vivo* transplantation of AKPT (*Apc^-/-^;Kras^G12D/+^;p53^-/-^;Tgfbr2^-/-^*) colon cancer organoids. Notably, we observed larger SOX2+ cell clusters in multiple regions of the tumour, suggesting clonal expansion of SOX2+ cells as the tumour progresses (Fig. 2d). Unlike in the premalignant tumour, these large SOX2+ clusters were not only SCA1+ but also KI67+ in the AKPT CRC (Fig 2e), indicating that SOX2+ cells proliferate and expand in advanced CRC. Previous studies have reported a dose-dependent role of SOX2 in retinal neural progenitor differentiation^17^ and pluripotency^36^. It is possible that the SOX2 expression level may influence cell cycle state in the tumour cells, where low SOX2 expression is slow-cycling and high SOX2 expression is proliferative.

To better understand the expression and cell state dynamics in the SOX2+ cells during tumour development, flow cytometry was performed with the AKPT tumours at Day 10 and Day 20 post transplantation (Fig. S2e). This revealed that the percentage of SOX2+ cells remained largely unchanged (∼8.1-8.5% of the total cancer cells) between the two time points, indicating that this cell population expands steadily as the tumour grows (Fig. S2f-g). On the other hand, cell state characterisation of this SOX2+ cell population showed a dynamic transition from the canonical LGR5+ cancer stem cell to SCA1+ fetal cell states between Day 10 and Day 20 (LGR5+SCA1-: 4.82% to 0.43%, P=0.502 ; LGR5+SCA1+: 14.5% to 9.55%, P=0.0008; LGR5-SCA1+: 72.3% to 89.4%, P<0.0001; respectively) (Fig. 2f). This suggests that expression of SOX2 reprogrammes LGR5+ canonical cancer stem cells to a fetal cell state over time in CRC (Fig. 2g).

Together, the results suggest that SOX2 expression is induced in a small number of cells soon after tumour initiation, but these remain as slow-cycling reprogrammed cells, possibly allowing them to evade immune surveillance. These rare SOX2+ cells undergo slow clonal expansion as the tumour develops, and eventually transition from slow-cycling to a proliferative state in advanced CRC, which expands the reprogrammed cell population and increase tumour plasticity.

### Ectopic expression of SOX2 induces acute intestinal hyperplasia

To examine the effect of SOX2 expression on cell state transition, we generated *Sox9iCreERT2; R26^Sox2/Sox2^* (Sox2 HOM) mice, which were used to induce SOX2 expression in the SOX9-expressing ISCs at the crypt (Fig. 3a). Strikingly, Sox2 HOM mice lost their body weight rapidly shortly after three consecutive oral gavage doses of tamoxifen administration (Fig. 3b and Fig. S3a). Small intestines harvested from the Sox2 HOM animals were significantly shorter than those from the control (*Sox9iCreERT2*) with visible inflammation in the proximal region (Fig. S3b-c). Histological analysis showed extensive intestinal hyperplasia of the Sox2 HOM animals with expansion of the crypt compartment (Fig. 3c). Expression of SOX2 was detected throughout the intestinal epithelium and was accompanied by high KI67 and CyclinD1 expression, indicating hyperproliferation of the SOX2-expressing cells (Fig. 3c and Fig. S3d). Interestingly, a marked reduction of SOX9 expression was noted in the transgenic crypts, suggesting its repression mediated by SOX2 (Fig. 1C). This is consistent with the SOX2-SOX9 expression dynamic previously described in lung cancer cells^37,38^.

**Figure 3.**
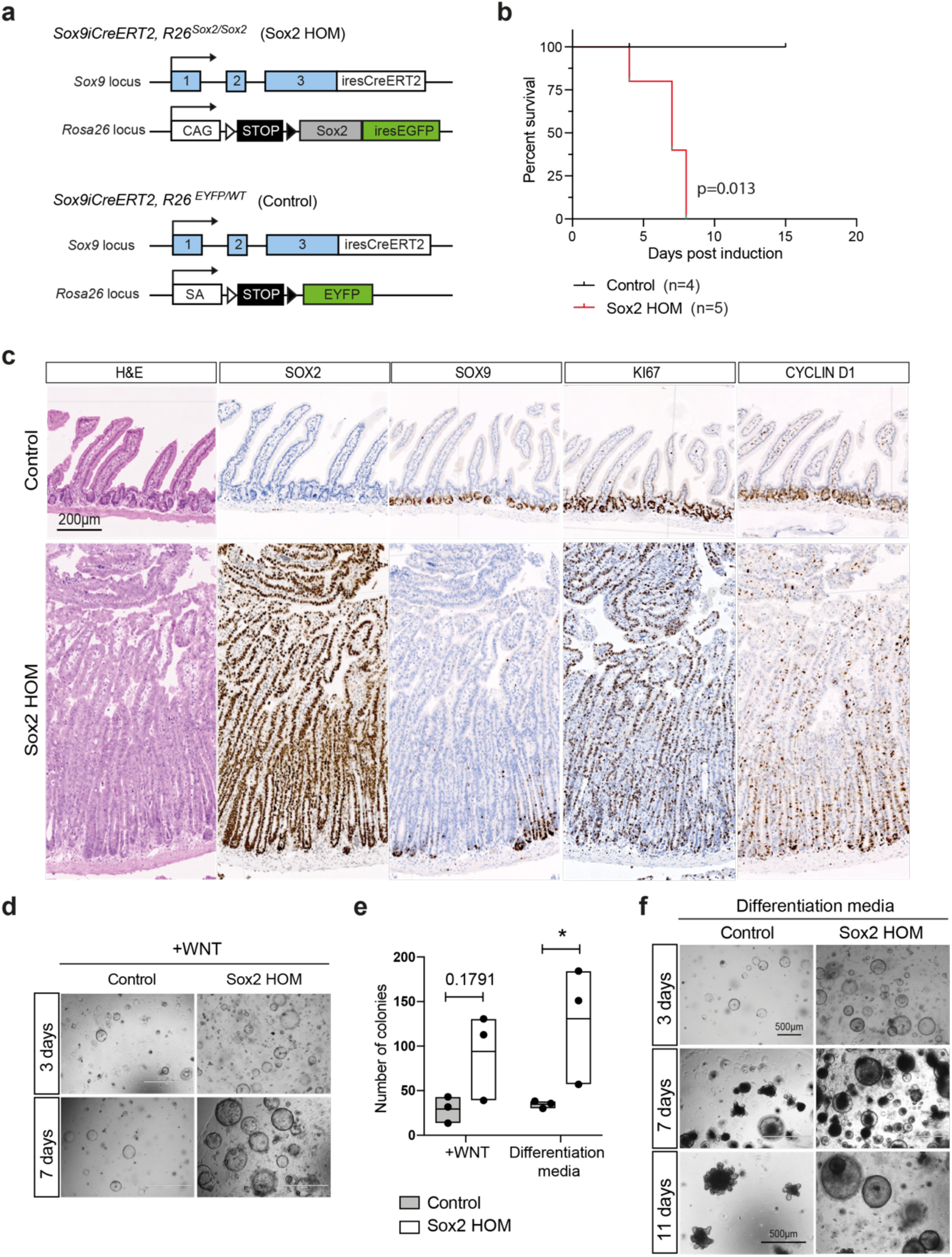
Ectopic expression of SOX2 induces acute intestinal hyperplasia. **a,** A schematic description of the *Sox9*and *Rosa26* locus at *Sox9iCreERT2/+*;*R26^Sox2-GFP/Sox2-GFP^* or *Sox9iCreERT2/+;R26^EYFP/WT^*allele targeting scheme. **b,** Kaplan-Meier survival analysis of *Sox9iCreERT2/+* (control, n=4) and *Sox9iCreERT2/+;R26^Sox2-GFP/Sox2-GFP^*(Sox2 HOM, n=5). p-values were determined using the Mantel-Cox test (∗P < 0.05). **c,** Representative histology (H&E) and immunohistochemistry (IHC) for SOX2, SOX9, KI67 and CYCLIN D1 of the small intestines of control and Sox2 HOM mice 7days after TM induction. Scale bar, 200μm. **d,f,** Representative images of organoids derived from control and Sox2 HOM intestines at 3, 7 or 11 days post-isolation for a colony-formation assay in WNT media (**d**) or differentiation media (**f**). **e,** Quantification of the number of colonies that form at day 7 after seeding 200 crypts in WNT media or differentiation media. Data represent mean ± s.e.m. of 3 biological replicates. *P<0.05, **P<0.01, ***P<0.001, 2-way ANOVA and Šídák’s multiple comparisons test.

We further explored the functional implication of SOX2-induced hyperplasia by evaluating the Sox2 HOM intestinal epithelial cells using *ex vivo* intestinal organoid culture^39^. Intestinal organoids derived from the Sox2 HOM animals formed more colonies and bigger spheroids compared to the control organoids in both WNT-supplemented (Fig. 3d-e) or differentiation media (Fig. 3e-f). WNT withdrawal induced differentiation in control organoids with classic budding morphology, whereas Sox2 HOM organoids remained as large spheroids, indicative of impaired differentiation by SOX2 expression (Fig. 3f).

### SOX2 expression inhibits intestinal differentiation and induces fetal reprogramming

To better understand the global impact of SOX2 overexpression, bulk RNA-seq was performed on the control and Sox2 HOM intestines. Previous studies have shown that ectopic expression of SOX2 in distal developing gut induces gastric-like reprogramming^24^. To evaluate if SOX2 induces gastric lineage specification in adult intestine, we included stomach tissues isolated from the control animals for comparison. Principal-component (PC) analysis of the expression levels of the Sox2 HOM intestine, control intestine and control stomach showed distinct clustering of the three sample types (Fig. S4a). The major variance across samples was attributed to the ectopic SOX2 expression in the intestines, whereas the control stomach clustered distinctively from the Sox2 HOM and control intestine (Fig. S4a). This indicates that SOX2 expression in the adult intestine did not cause major redirection to gastric specification. We therefore focused on the comparison between control and Sox2 HOM intestine in the subsequent analysis. Volcano plots revealed remarkably high numbers of differentially expressed genes (DEGs) (8037) between control and Sox2 HOM intestine (Fig. 4a, Supplementary Table 1), indicating that expression of SOX2 in the crypts induces extensive transcriptional reprogramming of the intestinal epithelium. Reassuringly, *Sox2* and its target *Sox21* were amongst the most upregulated DEGs. On the other hand, expression of *Sox9* was downregulated in Sox2 HOM intestines, in line with the protein expression observed earlier. We also observed downregulation of several key intestinal differentiation markers, including *Cdx2*, *Atoh1*, *Hnf1a*, *Defa2* and *Alpi*, suggesting that differentiation is inhibited by SOX2 expression. Ingenuity Pathway Analysis (IPA) showed that the S100 family, LPS/IL−1 Mediated Inhibition of RXR Function, tumour microenvironment and colorectal cancer metastasis pathways were amongst the most significantly enriched in the Sox2 HOM intestines (Fig. 4b). This correlates with the robustly activated upstream transcription factors *Yap1*, *Smad3-4*, *Fos*, *Jun*, nuclear factor kB (*NF-kB*) and various cytokines such as *Il6*, *Il1B*, *Tnf* and *Tgfb1* (Fig. S4b).

**Figure 4.**
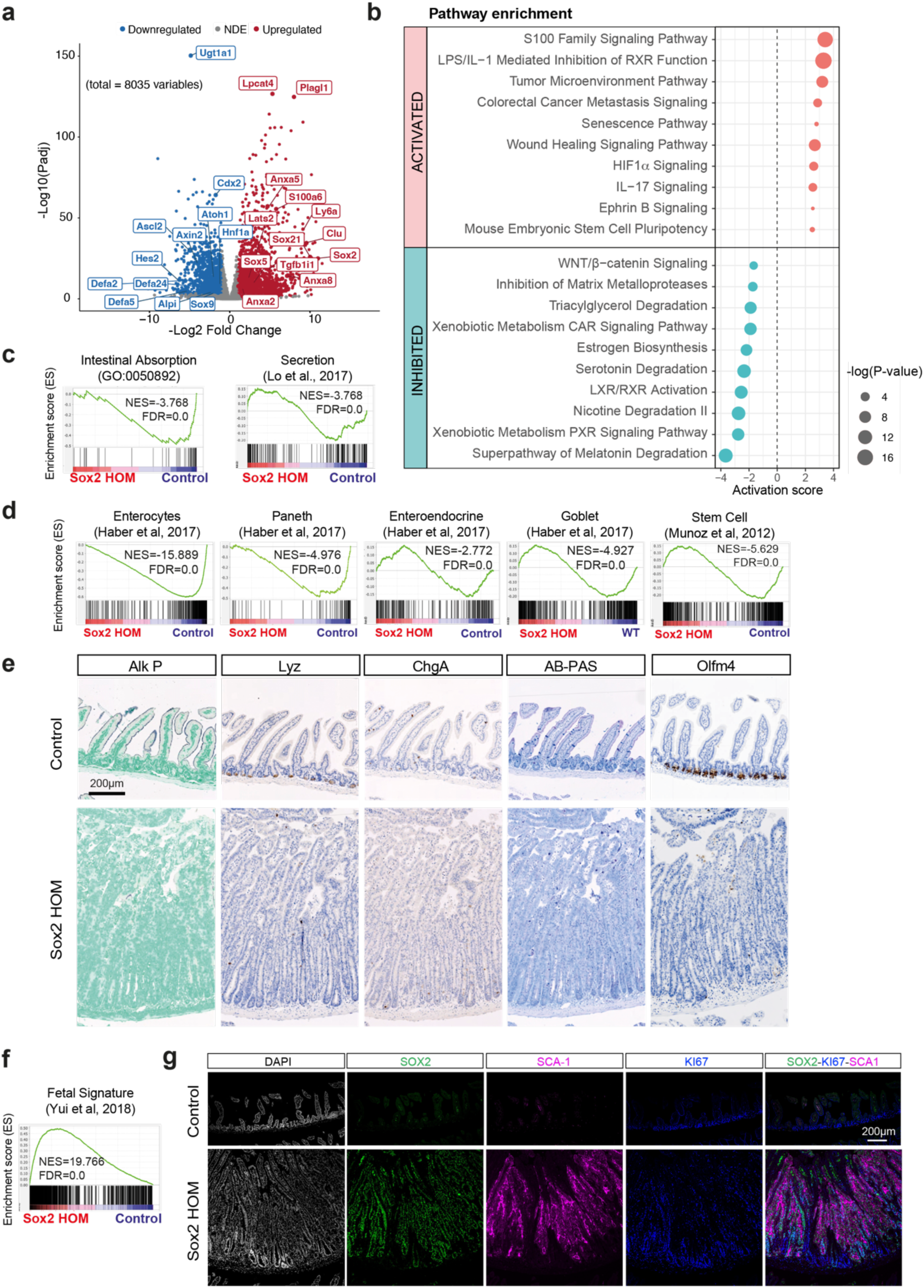
SOX2 expression inhibits intestinal differentiation and induces fetal reprogramming. **a,** Volcano plot showing significantly upregulated (red) or downregulated (blue) differentially expressed genes in control vs Sox2 HOM isolated crypts RNAseq. **b,** IPA Pathway Enrichment analysis showing activation score and P values of top 10 activated and inhibited pathways. **c,** Gene Set Enrichment Analysis (GSEA) for Intestinal Absorption and Secretion gene signatures in Sox2 HOM vs control intestine. **d,** GSEA of signatures related to stem cell presence and intestinal lineage differentiation (stem cell, enterocytes, Paneth, enteroendocrine and goblet gene signatures) in control vs Sox2 HOM isolated crypts RNAseq. **e,** OLFM4 (stem cell marker), Lysozyme (Paneth cell marker), CHGA (enteroendocrine cell marker) IHC and alkaline phosphatase (Alk P, enterocyte marker) and AB-PAS stainings (Goblet cell marker) in control and Sox2 HOM intestinal tissues. Images are representative of at least 4 animals analysed per group. Scale bar, 200μm. **f,** GSEA of fetal signature. **g,** Triple immunofluorescence for SOX2, SCA1 and Ki67 in control and Sox2 HOM intestines 7days after TM induction. Scale bar, 200μm.

Conversely, we observed downregulation of multiple metabolic pathways in the Sox2 HOM intestine (Fig. 4b), which could be attributed to loss of enterocyte differentiation. Indeed, many master regulators of intestinal epithelial differentiation such as the CDX family, FOXA family, SPDEF, HNF1 and the HNF4 family of transcription factors were strongly inactivated in the Sox2 HOM intestines (Fig. S4b). Consistently, gene set enrichment analysis (GSEA) showed inhibition of both absorption and secretion signatures in the Sox2 HOM intestine (Fig. 4c), which was confirmed by suppression of the individual enterocyte, Paneth, enteroendocrine and goblet cell lineage signatures (Fig. 4d). Immunostaining of Alkaline Phosphatase (enterocytes), Lysozyme (Paneth cells), Chromogranin A (enteroendocrine cells) and AB-PAS (goblet cells) further validated the loss of differentiation to all cell lineages in the Sox2 HOM intestines (Fig. 4e).

The lack of differentiation in the Sox2 HOM intestine prompts us to speculate if the hyperplastic crypts are caused by stem cell expansion. Surprisingly, GSEA showed a dramatic reduction of the ISC signature^40^ in the SOX2-expressing intestine (Fig. 4d), which is supported by the loss of ISC markers OLFM4 (Fig. 4e) and SOX9 expression (Fig. 3c) and a negative Z-score of WNT signalling pathway in the Sox2 HOM tissues (Fig. 4b). Interestingly, the mouse embryonic stem cell pluripotency pathway was activated in the Sox2 HOM intestine (Fig. 4b), suggesting fetal reprogramming in the tissues. Indeed, a remarkable enrichment of fetal gene signatures^5^ were observed in Sox2 HOM intestines (Fig. 4f), which was accompanied by the robust enrichment of the Yap signature^41^ and the nuclear translocation of YAP proteins (Fig. S4c-d). Enrichment of the wound healing signalling pathway and the regenerative revival stem cell signature^2^ was also noted in the Sox2 HOM intestine (Fig. 4b and S4c), suggesting that SOX2 expression in adult intestine activates tissue regeneration and reprogrammes the epithelial cells to a primitive state. Immunostaining of the fetal marker SCA1 confirmed its aberrant expression in the Sox2 HOM intestine, which was completely absent in the control tissue (Fig. 4g). Flow cytometry and immunostaining of the *ex vivo* organoids further demonstrated that SCA1 expression could be induced by WNT in control organoids, whereas ectopic expression of SOX2 could activate SCA1 in the absence of WNT (Fig. S4e-g). Of note, cleaved Casp3 staining was detected in the Sox2 HOM intestine, accompanied by enrichment of apoptosis and autophagy processes (Fig. S4h-i), suggesting that extensive epithelial reprogramming mediated by SOX2 may trigger cellular apoptosis and/or autophagy. Taken together, the data support the notion that SOX2 expression in adult intestine induces cellular plasticity by inhibiting differentiation and driving fetal reprogramming.

### Monoallelic SOX2 expression induces dormancy in young intestine but drives dysplasia in old intestine

Since SOX2 expression levels may influence cell cycle state^17,36^, we further evaluated the dose-dependent SOX2 expression in the intestine by generating heterozygous *Sox9^iCreERT2^*;*Rosa26^Sox2/+^*(Sox2 HET) mice to induce monoallelic SOX2 expression. A single low dose of tamoxifen was administered intraperitoneally in the Sox2 HET and Sox2 HOM young animals (12-15 weeks) to compare the dose-dependent effects of SOX2 expression in the intestine (Fig. S5a). Consistent with the earlier observation, Sox2 HOM animals lost weight soon after tamoxifen induction, with 4 out of 5 animals reaching the humane end point in less than 50 days (1 culled at 75 dpi) (Fig. S5b). On the other hand, 58% (11 out of 19) of Sox2 HET animals survived the induction (Fig. S5b). Of note, three of the Sox2 HET animals were culled for dermatitis, suggesting a role for SOX2 in the epidermis. In accordance with the high dose tamoxifen induction findings, histological analysis revealed dysplasia and aberrant crypt expansion in the Sox2 HOM intestine accompanied by hyperproliferation and loss of SOX9 expression. (Fig. 5a). In contrast, Sox2 HET intestine was largely intact with only sporadic expression of SOX2 in cells present in small clusters. To our surprise, these SOX2+ cell clusters were predominantly Ki67 negative (Fig. 5a). The results indicate that high SOX2 expression in adult intestine induces hyperproliferation, whilst low expression of SOX2 somehow triggers cell cycle arrest.

**Figure 5.**
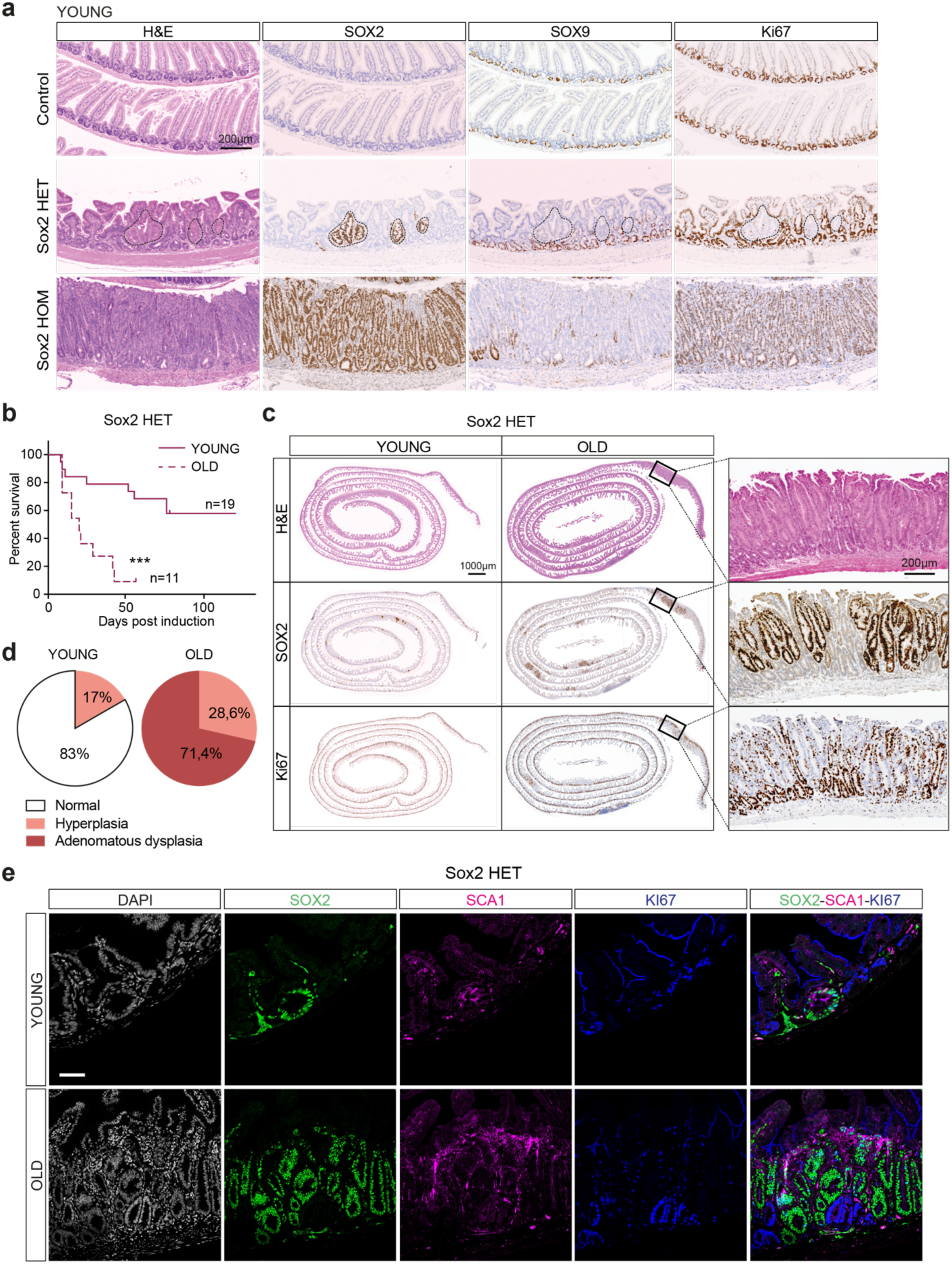
Monoallelic SOX2 expression induces dormancy in young intestine but drives dysplasia in old intestine. **a,** Representative H&E and IHC of SOX2, SOX9 and KI67 on intestinal sections from control, Sox2 HET and Sox2 HOM mice collected at the corresponding humane end points (Sox2 HET at 52dpi, Sox2 HOM at 9dpi) following 1 intraperitoneal (i.p.) dose of TM. Scale bar, 200μm. **b,** Kaplan-Meier survival analysis of Sox2 HET old and young animals following one i.p. dose of TM. P-values were determined using the Mantel-Cox test (∗∗∗P < 0.001). **c,** Representative H&E and IHC of SOX2 and KI67 on intestinal sections from Sox2 HET young (52dpi) and old (20dpi) mice following 1 TM i.p. dose. Scale bar, 1 mm for the full roll sections and 200μm for the enlarged image on the right. **d,** Pie chart showing the percentage of histopathological findings in the intestines of Sox2 HET young and old mice after tamoxifen induction. **e,** Representative IF of SOX2, SCA1 and KI67 on intestinal sections from Sox2 HET young (52dpi) and old (20dpi) mice following 1 TM i.p. dose. Scale bar, 200μm.

Since ageing increases cancer incidence^42^, we asked if tamoxifen induction in old Sox2 HET animals would accelerate tissue transformation. Single low dose tamoxifen was administered to both young (12-15 weeks) and old (55-70 weeks) Sox2 HET mice for comparison. Strikingly, the latter lost weight drastically soon after tamoxifen administration and all except one reached the preset humane endpoint by 43 days (Fig. 5b, S5c). Histopathological analysis showed tissue transformation of the intestine in all old Sox2 HET mice, with 71.4% of the animals developing adenomatous dysplasia and the rest showing tissue hyperplasia (Fig. 5c-d). In contrast, only 17% of the young Sox2 HET mice displayed tissue hyperplasia whereas the rest of the cohort had grossly normal intestines. The SOX2+ cell clusters in the old Sox2 HET intestine also appeared much larger in size compared to their young counterparts (Fig. 5c). Unlike the latter, immunostaining of the old tissues showed positive Ki67 expression in some SOX2+ cells (Fig. 5c), suggesting that SOX2-expressing cells proliferate and expand more efficiently in old Sox2 HET intestine. Of note, expression of Ki67 was much more prominent in the epithelium surrounding the SOX2+ clones (Fig. 5c), implying a non-cell-autonomous effect of SOX2 expression on the neighbouring cells. Furthermore, expression of SCA1 was detected in the SOX2+ clones in both young and old Sox2 HET intestine, suggesting that the SOX2-induced fetal reprogramming is not affected by cell cycle state (Fig. 5e, S5d). Although the SOX2+ clones were rare and slow-cycling in young Sox2 HET intestine collected at 52dpi (Fig. 5a), SOX2+KI67+ cells were readily detectable at an earlier time point (9dpi) (Fig. S5e). This implies that the initial SOX2 induction was efficient and proliferative in the Sox2 HET intestine, but most of the SOX2+ cells were subsequently eliminated from the intestine and the surviving small SOX2+ cell clusters are mostly in a dormant state. Indeed, strong macrophage infiltration was detected in both Sox2 HOM and HET intestine at an early time point (Fig. S5f-g), suggesting immune surveillance associated with the transgene to eliminate the SOX2- expressing cells.

Altogether, the results imply that abrupt SOX2 expression in the intestinal epithelium induces fetal reprogramming, which then triggers cell elimination by the host immune system. On the other hand, some SOX2+ cells may induce cell cycle arrest and thereby evade immune surveillance and remain as small cell clusters. It is conceivable that such an elimination process is partially compromised in ageing intestine, allowing clonal expansion of the SOX2+ cells and driving adenomatous dysplasia in the old Sox2 HET mice.

### SOX2 triggers reversible and p53-dependent senescence

One of the processes contributing to impaired cell clearance in ageing tissues is senescence. Since the senescence pathway was activated in the Sox2 HOM intestine (Fig. 4b), we investigated if expression of SOX2 induces senescence. First, we performed GSEA for the senescence signatures obtained from six different published datasets^43–48^. Significant enrichment of all the senescence signatures as well as the Senescence Associated Secretory Phenotype (SASP) signature^49^ was observed in the Sox2 HOM intestine (Fig. 6a-b). This was supported by the activation of several SASP-specific secreted factors such as *Hgf*, *Egf*, *Cxcl12*, *Gas6* in the Sox2 HOM intestine (Fig. S4b)^49^. Next, we explored if senescence is also induced in the Sox2 HET intestine by examining expression of the key senescence markers p16 and p21^50,51^. Immunostaining revealed both, with moderate levels in the SOX2+ clones in young animals and far more prominent staining in the adenomatous regions of old Sox2 HET mice (Fig. 6c).

**Figure 6.**
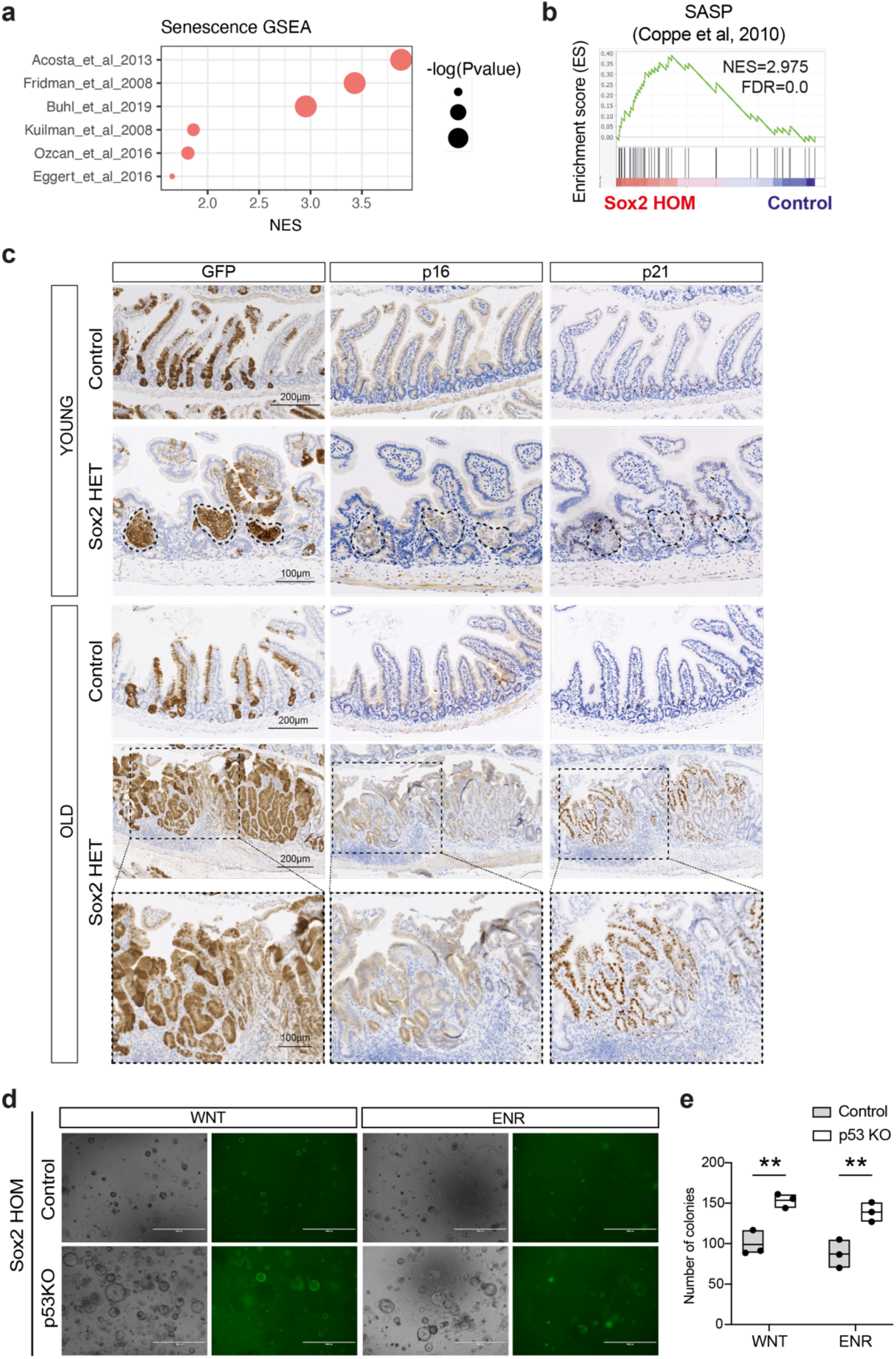
SOX2 triggers reversible and p53-dependent senescence. **a,** GSEA probing Senescence signature genes from the indicated publications in Sox2 HOM vs control intestinal crypts RNAseq. **b,** GSEA showing SASP signature enrichment in Sox2 HOM vs control intestinal crypts RNAseq. **c,** IHC using an antibody against GFP, p16 and p21 on intestinal sections of control and Sox2 HET young and old mice. Scale bars as indicated. **d,** Representative bright field (left) and GFP (right) images from a colony formation assay performed with control (Cas9)-Sox2 HOM and p53KO-Sox2 HOM intestinal organoids. Scale bar, 1000μm. **e,** Quantification of the number of colonies that form at day 7 after seeding 200 crypts in WNT media or ENR media. Data represent mean ± s.e.m. of 3 independent experiments **P<0.01, 2-way ANOVA and Šídák’s multiple comparisons test.

To functionally validate SOX2-induced senescence, organoids were derived from control and Sox2 HOM intestine and their growth capabilities were evaluated in different culture conditions. We showed earlier that the freshly derived Sox2 HOM organoids failed to differentiate upon WNT withdrawal from the media (Fig. 3f). Whilst organoids from both genotypes were able to grow efficiently with a cystic morphology in the presence of WNT, Sox2 HOM organoids underwent growth arrest soon after passaging in WNT withdrawal conditions (ENR) and remained as small, densely packed cell clusters (Fig. S6a). This became more striking in R-spondin withdrawal conditions (EN). Control organoids could not survive in the absence of both WNT and R-spondin (complete absence of extrinsic WNT signals), whilst the Sox2 HOM organoids continued to survive as small cell clusters (Fig. S6b). Despite being resistant to growth factor withdrawal, Sox2 HOM organoids were unable to expand and remained as dense cell clusters after passaging, indicative of cell cycle arrest. Indeed, strong β-galactosidase staining was detected in the Sox2 HOM organoids, suggesting that the cells were in a senescent state (Fig. S6c). The SOX2-induced senescence may explain the observation of SOX2+KI67- clones in early non-malignant adenomas. Given that SOX2+ clones can proliferate and expand in advanced CRC, we asked if loss of p53 may potentially reawaken senescent cells^52^. Remarkably, CRISPR targeting of p53 (p53KO) reactivated growth of the Sox2 HOM organoids (Fig. 6d and S6d), with both the size and number increased compared to controls (Fig. 6e and S6e). β-galactosidase staining was also reduced in the Sox2 HOM p53KO organoids (Fig. S6f), suggesting that SOX2-induced senescence is reversible and is dependent on p53.

Taken together, these results indicate that SOX2-induced reprogramming triggers senescence-mediated dormancy^53^, licensing the cells to evade surveillance^45,54^. Most of the senescent SOX2+ cells will be eliminated through immune clearance, leaving behind small clusters of rarely cycling SOX2+ clones survived in the early non-malignant adenomas. As tumorigenesis progresses, oncogenic mutations (such as p53 inactivation) may reverse senescence, leading to expansion of SOX2+ reprogrammed cells observed in advanced CRC.

### SOX2-induced fetal reprogramming sustains drug tolerance

Since fetal reprogramming and dormancy have been associated with drug-tolerant persisters (DTP)^55^, we asked if SOX2 contributes to the DTP state in CRC. Treatment of WT and Sox2 HOM organoids with chemotherapy drug 5-Fluorouracil (5FU) showed increased drug tolerance from IC50 4.7mM±1.22 in WT to IC50 13.8mM±2.09 in Sox2 HOM (Fig. 7a), suggesting that SOX2 expression promotes drug tolerance. Since AKPT tumours contained SOX2+ cell clusters, we asked if SOX2 contributes to chemoresistance in advanced CRC. Treatment of AKPT organoids with high doses of 5FU (75mM and 125mM) caused pronounced cell death (Fig. S7a). The residual persister cells showed upregulated expression of SOX2 in the 5-FU treated AKPT organoids cultured in both 2D and 3D conditions (Fig. 7b and S7b). To test if SOX2 contributes to drug tolerance in AKPT CRC organoids, we deleted *Sox2 b*y CRISPR targeting (Fig. S7c). This caused a drastic change of morphology of the AKPT organoids from the typical large cysts to irregular folding structures (Fig. S7c). To our surprise, these SOX2 KO organoids seemed to grow faster than the parental controls with increased cell viability (Fig. S7d). Importantly, loss of SOX2 resulted in robust reduction of reprogramming marker expression at both transcription (Fig. 7c) and protein levels (Fig. 7d), indicating that SOX2 is required to sustain a fetal-like cell state in AKPT tumour cells. We further assessed if inhibition of SOX2 and the fetal cell state would impact drug response in AKPT organoids. Treatment with 5-FU upregulated *Sox2* expression in parental AKPT organoids but, as expected, not in the two isogenic SOX2 KO counterparts (Fig. S7e). Viability assays further showed that the IC50 decreased from 40.88mM±29.12 in AKPT control to 1.78mM±0.88 and 1.57mM±0.44 in the SOX2 KO-1 and KO-2 respectively (Fig. 7e). Overall, the data support the notion that the cystic morphology of parental AKPT organoids is indicative of a fetal-like state, whereas *Sox2* deletion suppresses fetal reprogramming in the advanced CRC organoids, leading to loss of spheroid appearance and decreased drug tolerance (Fig. 7f).

**Figure 7.**
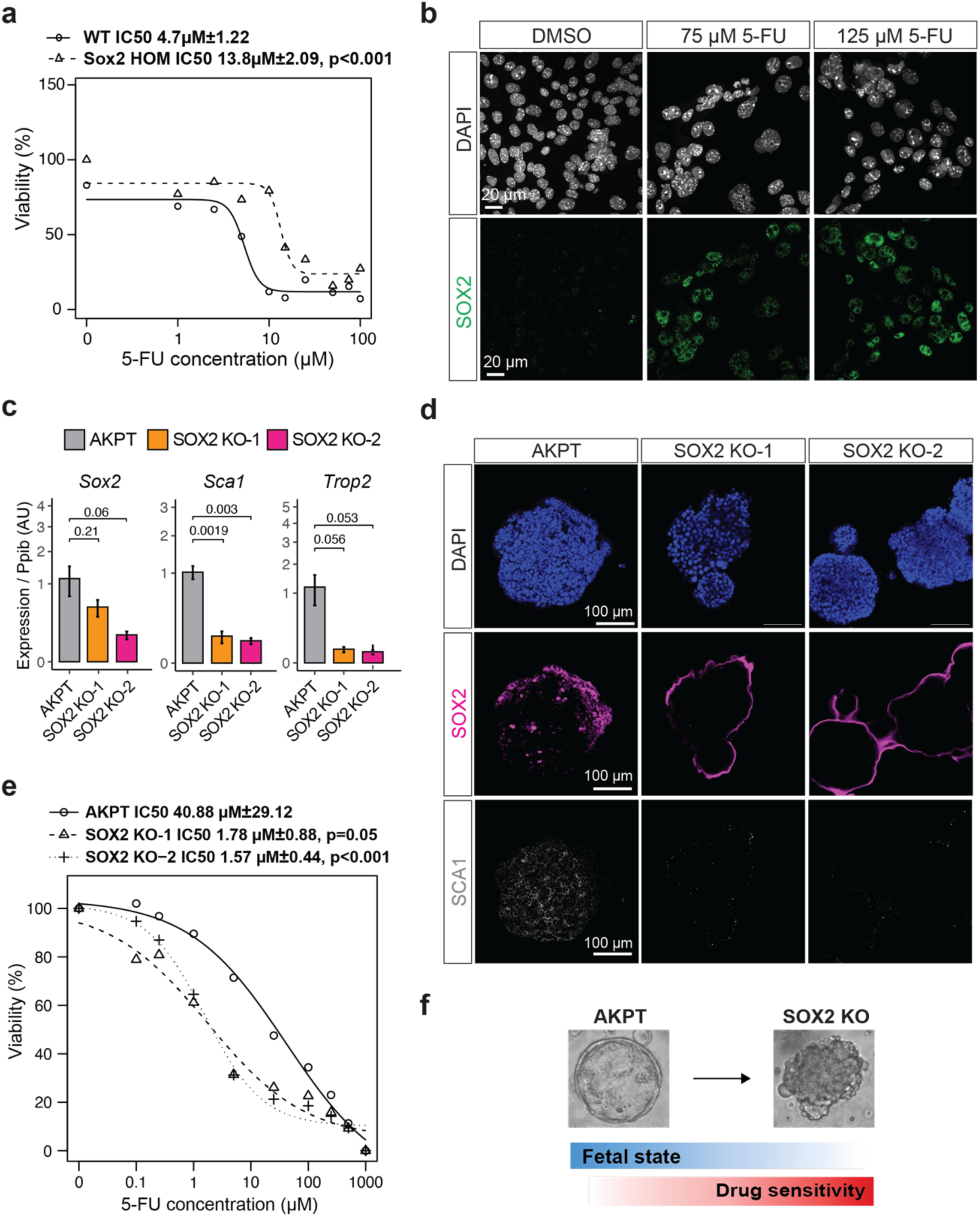
SOX2-induced fetal reprogramming sustains drug tolerance. **a,** 5FU dose-response curves for Sox2 WT and Sox2 HOM intestinal organoids. The half maximal inhibitory concentration (IC50) for each of the organoid lines was calculated from the viability measurements at different 5FU concentrations using a model fitting function. The model is generated with the averaged value of four technical replicates, in four biological replicates. The IC50, deviation and p values are included in the top of the plot. **b**, Representative confocal immunofluorescence of Sox2 and DAPI of the AKPT cell line treated with 5-FU at 75 µM, 125 µM, and DMSO control. Scale bar: 20 µm. **c,** qPCR quantification of *Sox2*, *Sca1*, and *Trop2* in Sox2 WT and two Sox2 KO models from AKPT organoids. Expression values are normalized to Ppib levels and represented as Log2 Fold Change. The experiment was performed in three biological replicates, and each sample was measured in three technical replicates for the qPCR quantification. Welch Two Sample t-test was used to test the differences between the WT and the two KO organoid models. **d**, Representative immunofluorescence of SOX2 and SCA1 in Sox2 WT and two Sox2 KO AKPT organoids. Scale bar: 100 µM. **e**, 5FU dose-response curves for WT and two Sox2 KO models from the VAKPT cell line. IC50 for each of the cell lines was calculated from the viability measurements at different 5FU concentrations using a model fitting function. The model is generated with the averaged value of four technical replicates, in three biological replicates. The IC50, deviation and p values are included in the top of the plot. **f**, Schematic summary showing loss of SOX2 in AKPT organoids inhibits cystic morphology and fetal cell state, which in turn sensitises chemotherapy response.

## Discussion

Tumour plasticity has been linked to cancer metastasis and response to therapy^1^. It is believed that some tumour cells adopt a reversible slow-cycling cell state, a phenotypic switch that allows them to survive host immune cell surveillance and drug treatment. Despite extensive efforts, the molecular drivers of tumour plasticity remain elusive. SOX2 has been associated previously with tumour plasticity in glioblastoma and prostate cancer^56,57^, but these studies focus on lineage dedifferentiation to cancer stem cells or progenitor cells rather than fetal reprogramming. Thus far, most studies report that SOX2 expression in cancer cells drives cell proliferation^58^. In this study, we show that SOX2 induces fetal reprogramming and reversible senescence-induced dormancy in CRC. SOX2 is upregulated in human CRC and correlates with poor prognosis. We find that SOX2 marks a reversible, slow-cycling and plastic subpopulation of CRC cells. The slow-cycling SOX2+ cells are first detected in rare small cell clusters of *APC*-mutated early adenomas that are largely dormant. Expression analysis and transgenic mouse models demonstrate that SOX2 drives fetal reprogramming in normal intestinal epithelium and in cancer. SOX21, a downstream target of SOX2, has been previously reported to regulate pluripotency^31^. Our data indicate that SOX21 is induced in SOX2- expressing intestine and is also enriched in PDS3 CRC, suggesting that both members of the SoxB gene family coordinate together to drive reprogramming in CRC. It has been shown that inducing pluripotency with the four reprogramming factors (Oct4, Sox2, Klf4, and c-Myc) triggers senescence^53^. Consistently, we demonstrate that SOX2 alone is sufficient to induce reprogramming and senescence in the intestine, licensing the SOX2+ cells to evade immune surveillance and cell clearance triggered by reprogramming^59^. This results in small slow-cycling SOX2+ cell clusters detected in early non-malignant tumours. Subsequently, p53 loss, which is seen in advanced CRC, reverses senescence and reawakens the SOX2+ cells, which then undergo clonal expansion and promote tumour plasticity. Loss of SOX2 in AKPT organoids robustly inhibits expression of fetal markers and accelerates tumour cell growth, suggesting that its expression is crucial in sustaining fetal reprogramming and a senescence-induced dormant state. This is potentially important for promoting metastasis by driving tumour cells to a dormant/senescent state able to escape surveillance and survive a harsh environment in the absence of a surrounding niche during dissemination. It will be important to further explore the potential role of SOX2 in CRC metastasis. In particular, it would be interesting to test if SOX2+ cells are the origin of micrometastases at secondary sites.

Besides metastasis, a dormant and plastic state also plays a crucial role in drug tolerance and cancer relapse. Cancer cells possess the ability to acquire a reversible DTP state, which allows them to evade therapeutic pressures such as chemotherapy^1,55^. Our data show that SOX2 is upregulated in the residual persister cells following chemotherapy treatment, suggesting that SOX2 marks DTP cells in CRC. Loss of function analysis further demonstrates that SOX2 is required for sustaining a fetal-like state, indicating its functional role in drug tolerance. Given that this rare slow-cycling subpopulation is detected in early adenomas, SOX2+ cells may mark pre-existing primed DTP cells and help provide intrinsic tolerance to drugs. Therapeutic pressure may further activate SOX2 expression to acquire a DTP state, which in turn may allow the tumour cells to adopt a reversible senescent state and consequently to evade cell death. Understanding upstream drivers of SOX2 expression will be important for targeting the DTP state. Previous studies have suggested approaches for DTP targeting, such as inhibiting repressed chromatin, PI3K–mTOR, TGFb or YAP/TAZ signalling^1^. Given that SOX2 is absent in normal intestine but marks the DTP state in cancer, this may represent an alternative tumour-specific DTP targeting strategy for CRC.

Our findings also reveal that SOX2 expression in intestinal epithelial cells in aged animals makes them more prone to transformation to adenomatous dysplasia compared to young animals. It is well known that senescence plays a significant role not only in cancer, but also in age-related diseases. Accumulation of senescent cells due to increased DNA damage, dysregulation of SASP and impaired immune clearance of senescent cells have been reported in ageing tissues^60^. For instance, SASP may intensify age-related disruption by extending the senescence phenotype to neighbouring cells through cytokine secretion^43^. The self-elimination programme by the body’s immune system may also be compromised in aged tissues^61^, resulting in inefficient removal of senescent cells. Reduced senescent cell clearance may explain the increased SOX2+ cells observed in ageing Sox2 HET mice, eventually leading to enhanced tumour transformation. Since sporadic *APC* mutations could happen years, if not decades, preceding tumour development and diagnosis^62^, it is conceivable that SOX2 activity may, at least in part, contribute to increased CRC rates in elderly populations. Further study of SOX2 may clarify its role in age-related cancer. Recent evidence suggests that senescence can be exploited to treat age-related diseases^63–65^ and cancer^66,67^. Future work on characterising the mechanisms underlying oncogene-induced senescence will help develop targeting strategies against senescence-induced DTPs in cancer.

## Acknowledgements

We thank the Francis Crick Institute’s Biological Research Facilities, Experimental Histopathology, Advanced Sequencing facilities, Bioinformatics and Biostatistics for technical support.

## Materials and Methods

### Patients and specimens

All colon cancer specimens were obtained from Basque BioBank upon informed consent and with evaluation and approval from the corresponding ethics committee from Basque country (CEIm Euskadi). A table providing clinic-pathological information is included as Supplementary Table 2.

### Animals and Drug Administration

All mouse experiments were performed in accordance with the guidelines of the Animal Care and Use Committee at the Francis Crick Institute. All animals were maintained with appropriate care according to the United Kingdom Animal Scientific Procedures Act 1986 and the ethics guidelines. All animal regulated procedures were carried out according to UK Home Office guidelines and regulations (PEF3478B3, PP4021677 and PP8826065). *Sox9^tm1(cre/ERT2)Haak^* (*Sox9^iresCreERT2^*)^1^, *Rosa26R-Sox2-IRES-GFP*^2^, *Lgr5-EGFP-IRES-CreERT2^3^* and Apc^fl/fl4^ mouse strains were used in this study. Mice were inbred on a C57BL/6J background and successive crosses were performed to combine alleles. Experiments were either performed using littermates or animals with comparable backgrounds. CreERT2 activity was induced by tamoxifen oral gavage administration for 3 consecutive days at 5mg/25g body weight/day (Fig 3 and S3) or one intraperitoneal injection at 1.5mg/10g mouse weight (from a 20mg/ml stock solution) (Fig 5 and S5). For proliferation analysis, animals were injected intraperitoneally (IP) with 30 mg/kg EdU (E10187, Molecular Probes) 2 hours before tissue collection. Mice were culled by schedule 1 procedure (S1K) at the desired time point or when losing 10% of their body weight at the time of the injection. Genotyping was performed by Transnetyx®.

### Establishment and maintenance of mouse organoid cultures

Organoids were established from freshly isolated adult small intestine, as previously described^8^. Briefly, 5cm of duodenal small intestinal tissue was opened longitudinally and villi were scraped using a cover slip. The remaining tissue was incubated in 15mM EDTA with 1.5mM DTT at 4°C for 10min and moved to 15mM EDTA solution at 37°C for an extra 10min. Subsequently, the tissue was shaken vigorously for 30sec to release epithelial cells from basement membrane and the remaining remnant intestinal tissue was removed. Cells were washed once with Advanced DMEM/F12 (Thermo Fisher, 12634010) containing Glutamax (Termo Fisher, 35050061), Hepes (Thermo Fisher, 15630056) and P/S (Penicillin-Streptomycin, Thermo Fisher, 15070063) filtered through a 70μm cell strainer and resuspended in Cultrex BME Type 2 RGF Pathclear (Amsbio, 3533-01002). All freshly isolated organoids were maintained in either Intesticult medium (Stem Cell technologies, #06005) or in-house made basal medium containing EGF (Invitrogen PMG8043), Noggin, Rspondin and Wnt3A WENR medium, as previously described^9^. The Rho kinase inhibitor Y-27632 (Sigma, Y0503) was added to the culture during first week of crypt isolation and single cell dissociation.

Noggin and R-Spondin conditioned media were generated by HEK293T cells. Wnt3A conditioned medium was generated from L cells. All images were acquired using an EVOS FL Cell Imaging System (Life technologies).

For organoid formation assay, crypts were counted using a brightfield microscope and 200 crypts were seeded in 20μl of Cultrex BME Type 2 RGF Pathclear in individual wells of a 48-well plate and cultured in WENR medium for 5 days until counted. Three technical replicates were performed per animal.

For R-Spondin withdrawal challenge, organoids were passaged and seeded in 3 x 10μl droplets per well of a 24-well plate. Organoids were allowed to re-establish in normal ENR medium (5% R-Spondin) for 48hr. Subsequently, organoid medium was replaced and organoids were cultured in either ENR or EN medium (without R-Spondin).

### Xenograft experiments

*Apc^-/-^;Kras^G12D/+^;p53^-/-^;Tgfbr2^-/-^*(AKPT) organoids were maintained in in-house made basal medium and expanded in Cultrex BME Type 2 RGF Pathclear. Organoid suspension was subcutaneously injected into both flanks of 8-week-old NSG mice. The xenografts were allowed to grow for a maximum of 20 days, with continuous monitoring of tumour size. Tumours were collected at either day 10 or 20, weighed, and harvested either for paraffin embedding or flowcytometry downstream protocols. All mice were housed in a pathogen-free environment and handled in strict accordance to institutional protocols.

### b-galactosidase staining for senescence detection

Intestinal organoids cultured in 8-well Nunc™ Lab-Tek™ II Chamber Slide™ System were washed once with PBS and fixed in SA-β-galactosidase (SA-β-gal) staining fix solution for 15 min at room temperature (Senescence β-Galactosidase Staining Kit #9860, CST). Cells were then washed three times with PBS and incubated with SA-β-gal staining solution following manufacturer instructions for 16–20 h at 37°C. 300mL of β-gal staining solution was added to each well and the plate was sealed with Parafilm to prevent evaporation of the staining medium. After the overnight incubation, cells were washed with PBS and observed under a bright field Leica DM500 microscope.

### Immunohistochemistry and immunofluorescence

For analysis of small intestine by immunohistochemistry (IHC), tissues were fixed in 10% formalin for 24h and embedded in paraffin. Sections were deparaffinized with xylene and rehydrated in a graded series of ethanol. Antigen-retrieval was performed for 20 min at ∼95-100°C in 0.01M Citrate (pH 6) or Tris-EDTA buffer (10mM Tris base, 1mM EDTA, pH 9). Slides were then treated using the relevant blocking buffer (10% Normal goat serum or 10% Normal donkey serum in 1% BSA) and incubated overnight with the chosen antibody at 4°C (see Supplementary Table 3 for full list of antibodies). Slides were washed in PBS and then incubated with the secondary antibody for 1h at room temperature and washed again with PBS before developing. For colorimetric staining with diaminobenzidine (DAB), slides were incubated with peroxidase substrate, dehydrated, counterstained with Hematoxylin solution according to Mayer (Sigma, 51275) and mounted. Slides were scanned using a Zeiss slide scanner and images were processed using QuPath^10^ and Zeiss ZEN Lite Software.

For immunofluorescence, slides were incubated with Alexa-Fluor 488, Alexa-Fluor 568 or Alexa-fluor 647 antibody for 1h following the primary antibody incubation, washed three times with PBS, incubated with 4’,6’-diamidino-2-phenylindole (DAPI) for 15 min to visualize nuclear DNA and mounted with ProLong Gold Antifade Mountant. Images were acquired as z-stacks using a Leica SPE or a Leica SP8 confocal microscope and processed using Fiji.

When indicated, sections were stained for Hematoxylin & Eosin (H&E), alkaline phosphatase (Alk P) and Alcian Blue-Periodic Acid Schiff (AB-PAS) staining. EdU was detected according to the manufacturer’s protocol (Thermo Fisher Scientific, Click-iT Plus EdU Alexa Fluor 555 imaging kit C 10638) to evaluate proliferating cell number.

For the analysis of the role of SOX2 in the response to chemotherapy, a 2D version of the AKPT mouse colorectal organoids were engineered with two sets of three guides targeting the *Sox2* locus (see CRISPR-Cas9 gene editing section for details). The resulting cells were seeded on un-coated glass slides in a 24-well plate at a confluence of 40,000 cells per well and incubated for 72 hrs to adhere. Cells were treated with 5-Fluorouracil (5-FU, F6627, Sigma-Aldrich) at the concentrations indicated or DMSO for an additional 72 hrs. At the endpoint, 4% PFA was used to fix the cells, followed by permeabilization with 0.1% Triton/PBS and blocking with 5% Normal Goat Serum in PBS 0.1% Triton. The samples were incubated with primary antibodies for SOX2 (Santa Cruz Biotechnology, sc-365823) and SCA1 (Santa Cruz Biotechnology, sc-52601) overnight. For secondary antibodies, Alexa-Fluor 488 or Alexa-Fluor 568 were used and incubated for 1h, then stained with DAPI for 10 min to visualize nuclear DNA, washed three times with PBS, and mounted with ProLong Gold Antifade Mountant. Images were acquired as z-stacks using a Leica SP8 confocal microscope and processed using Fiji.

Organoids were re-derived from the AKPT cell line by growing them in BME and organoid culture media. Once organoids were formed, they were seeded in an IBIDI 8-well chamber. They were left 24h to settle and then treated with DMSO or 250 µM 5-FU for 72 hrs. The organoids were then fixed with a 4% PFA solution, incubated with NH_4_Cl to quench autofluorescence, and permeabilized with 0.3% TritonX-100. To amplify the SOX2 signal in the AKPT organoids we used an adaptation of the Tyramide Signal Amplification system. Therefore, we incubated the sample for 30 min with a 0.3% H_2_O_2_ solution, blocked with 5% normal goat serum solution and stained with anti-SOX2 (Santa Cruz Biotechnology, sc-36582), or anti-Sca1 (Santa Cruz Biotechnology, sc-52601) overnight. For secondary antibodies except SOX2, Alexa-Fluor 488, Alexa-Fluor 568, or Alexa-Fluor 654 were used together with DAPI to visualize nuclear DNA. After the secondary incubation, we incubated the samples with the ImmPRESS HRP goat anti-mouse IgG Polymer kit (peroxidase) (Vector laboratories, MP-7452) for 30 min. The SOX2 fluorescent signal was detected using the Tyramide Signal Amplification Alexa-fluor 555 fluorophore (Thermo Fisher Scientific, B40955) and incubated for 10 min, washed three times with PBS, and mounted with ProLong Gold Antifade Mountant. Images were acquired using a Zeiss LSM 880 inverted confocal microscope with AiryScan.

### Flow cytometry

#### Organoids

Sox2-HOM and WT intestinal organoids were collected using cold Advanced DMEM/F12 (Thermo Fisher, 12634010) containing Glutamax (Termo Fisher, 35050061), Hepes (Thermo Fisher, 15630056) and P/S (Penicillin-Streptomycin, Thermo Fisher, 15070063), followed by one centrifugation wash in cold PBS. The pellets were incubated with TrypleE (Gibco, Waltham, MA; 12604013) for 10 minutes to dissociate to single cells. TrypLE was stopped by adding Advanced DMEM (Gibco; 12491015) containing 10% fetal bovine serum (FBS) (Gibco; 10270106) and dissociated cells were passed through a 20-μm strainer. Cells were centrifuged and resuspended in complete Advanced DMEM/F12 containing Rho kinase inhibitor Y-27632 (Sigma, Y0503) until they were used. Before staining, cells were resuspended in FACS Buffer FB, consisting of 1% BSA and 2mM EDTA, vortexed and transferred to a 96-well plate with conical shaped wells (Thermofisher, 277143). Samples were then incubated on ice in the dark with APC anti-mouse Ly-6A/E (Sca-1) Antibody (clone D7, 108111, BioLegend) antibody for 45 min, washed once with FB, resuspended and filtered through 70μm Filcons into 5ml polypropylene FACS tubes Flow cytometry was performed on a Fortessa BD FACSAria II System (BD Biosciences) using the gating strategy described in the manuscript. Graphs and data analysis were performed in FlowJo 10.10.0.

#### Tumour samples

Tumour samples were quickly transferred into ice-cold phosphate-buffered saline (PBS) (Fisher Scientific, 10091403) on ice. All non-tumour tissue that was attached to the outer edge of the denser tumour core was removed, the surface of the tumour samples was then carefully dried with sterile paper and tumour weight recorded. Samples were washed once with ice-cold RPMI media and minced with disposable scalpels in 2 mL of disaggregation buffer (DB), consisting of 2 mg/mL Collagenase Type IV (Thermo Fisher, 17104019), 1 mg/ml DNase1 (Sigma Aldrich, 10104159001) and 0.5 mg/ml Hyaluronidase (Sigma Aldrich, H3757) in RPMI. Tumour pieces less than 3mm in length were transferred to a C-tube (Miltenyi Biotech, 130-096-334) and a further 3ml of DB added. If tumours were >600 mg, the disaggregation was carried out in two C-tubes, each with a total of 5 mL DB. The C-tube was placed in a GentleMACS Octo Dissociator (Miltenyi Biotech), heating blocks fitted, and tumours disaggregated using the automated 37C_m_TDK1 program. Once complete, the C-tube was centrifuged at 100 g for 2 min to ensure the contents were gathered at the bottom of the tube.

The sample was diluted with a further 5ml of fresh and warmed DB, mixed well by pipetting and filtered into a 50ml tube through a 70 μm strainer, which was then washed with 10ml ice-cold RPMI to quench the digestion. The single cell suspension was pelleted at 300 g for 6 min and used for flow cytometry staining. Before starting with the staining, cells were resuspended in FACS Buffer FB, consisting of 1% BSA and 2mM EDTA, vortexed and transferred to a 96-well plate with conical shaped wells (Thermofisher, 277143). Samples were first incubated with Live/Dead Aqua staining solution (L34976, Invitrogen) in PBS for 15min at 4°C. After adding 100ul of PBS per well, the plate was spined 2min at 1700rpm and the supernatant was discarded. Samples were incubated in FB containing Fc block (1/100) for 30min at 4°C followed by washing step with FB. Primary antibodies were added for 45min at 4°C (see Supplementary Table 3) in the dark. Samples were washed with FB twice and supernatants were discarded. A 30min permeabilization step using True-Nuclear™ Transcription Factor Buffer Set (424401) was performed before staining for SOX2 (561610, BD Biosciences) during 60min. Following manufacturer’s instructions, cells were washed in permeabilization buffer and were finally resuspended and filtered through 70 μm Filcons into 5 mL polypropylene FACS tubes. Flow cytometry was performed on a LSRFortessa^TM^ X20 System (BD Biosciences) using the gating strategy described in the manuscript. Graphs and data analysis were performed in FlowJo 10.10.0.

### Viability curves

Organoids in culture were incubated for 5 minutes in TypLE at 37 °C to dissociate them into single cells. 2,500 cells were seeded in 10µl of BME in each round bottomed well of a 96 well plate. Three days after seeding, nine increasing 5-FU doses (0.1, 0.25, 1, 5, 25, 100, 250, 500, 100µM) and DMSO as control were added to the plates. After three additional days of incubation, CellTiter-Glo 3D Cell Viability Assay (Promega, G9681) was used to determine the viability of the organoids following manufacturer instructions. The assay was transferred to a flat clear-bottom plate with white opaque wells for the measurements and the luminescent signal was determined using a TECAN Spark Microplate Reader. All treatments were performed with four technical replicates and each assay was performed in three biological replicates. The data was normalised, processed and represented using the R package “drc, Analysis of Dose-Response Curves” version 3.0-1.

### Immunoblotting

Organoids were lysed in cold lysis buffer containing 150mM NaCl, 30mM Tris (pH 7.5), 1mM EDTA, 1% Triton X-100, 10% Glycerol, 0.5mM DTT and Halt™ Protease and Phosphatase Inhibitor Cocktail (Thermofischer, 78440). Lysates were centrifuged for 30 min at 15000 rpm at 4°C and supernatants were kept for downstream protein quantification using Thermo Scientific™ Pierce™ BCA Protein Assay Kit (Thermofischer, 10741395) according to manufacturer’s instructions. Upon quantification, equal amounts of protein were mixed in 4x Laemmli Sample Buffer (Biorad, 1610747), resolved in 10% sodium dodecyl sulfate-polyacrylamide gels (SDS-PAGE), and subsequently transferred to polyvinylidene difluoride (PVDF) membranes. Membranes were blocked using 5% milk in Tris-buffered saline (50mM Tris, 150mM NaCl, pH 7.6) containing 0.1% Tween-20 (TBS-T) for 1hr and anti-p53 (DO-1) (Santa Cruz, sc-126), was added to blocking solution. Primary antibody incubations were carried out at 4°C overnight. After washing with TBS-T, the appropriate HRP-conjugated antibody was added for 1hr at room temperature. Anti-β-actin-HRP (Sigma, A3854) was used as a loading control. Membrane was then washed, and antibodies were detected using GE Healthcare Amersham™ ECL Prime Western Blotting Detection Reagent (GE Healthcare, 12316992). Protein bands were visualised using an Amersham Imager 680.

### Bulk RNA sequencing sample preparation

RNA was extracted from dissected male and female proximal intestines and stomach using the RNeasy Micro kit (Qiagen) according to the manufacturer’s protocol. Harvested tissue was suspended in RLT buffer (included in the kit) supplemented with 40mM Dithiothreitol (DTT) to inhibit RNases activity. To enhance the yield of extracted RNA, the samples underwent one freeze/thaw cycle prior to the RNA extraction step. RNA integrity (RIN) was examined using Bioanalyzer 2100 RNA 6000 Nano kit from Agilent and RIN cut-off was set to 7. Libraries were prepared using KAPA mRNA HyperPrep kit (KK8580) according to manufacturer’s instructions. Libraries were quantified using the TapeStation (Agilent) and pooled in equimolar proportions. Libraries were sequenced on an Hiseq4000 (Illumina), to achieve an average of 25 million reads per sample. The sequencing was performed on biological triplicates for each data point.

### RNA-Seq data analysis

Fastq files were processed using the nf-core/RNASeq pipeline (10.5281/zenodo.4323183) version 3.0 using the corresponding versions of STAR RSEM to quantify the reads against release 95 of Ensembl GRCm38.89 version transcriptome. All parameters for RSEM were run as default except “–forward-prob” which was set to 0.5. These raw counts were then imported into R (R Core Team (2020). R: A language and environment for statistical computing. R Foundation for Statistical Computing, Vienna, Austria. URL https://www.R-project.org/) version 4.03 /Bioconductor version 3.12^11^. Differential expression analysis (DE) was performed using DESeq2 ^12^version 1.30.1. Differentially expressed genes were defined as those showing statistically significant differences between pairwise groups if the adjusted P value was less than 0.05 (FDR < 0.05).

Gene set enrichment analysis (GSEA) was performed using the GSEA desktop software (version 4.1.0) using the following parameters: GSEA Preranked > no collapse of gene symbols > classic enrichment statistic > Chip platform “Mouse_Gene_Symbol_Remapping_Human_Orthologs_MSigDB.v7.5.chip”. GSEA custom lists were obtained from the indicated publications. For Metacore analysis (https://portal.genego.com/) and Qiagen Ingenuity Pathway Analysis (IPA), gene lists of upregulated and downregulated genes were created by using FDR<0.05 and FC>1.5 cut-offs. One click analysis included Pathway Maps and GO Processes for Metacore and Upstream regulators and Canonical Pathways enrichment for IPA.

### TCGA data analysis

TCGA Pan-Cancer (PANCAN) batch effect normalized gene expression RNA-Seq data and the corresponding clinical information for colon and rectal adenocarcinoma (n = 607) and adjacent normal tissue (n = 51) were obtained from Xenahubs using the “UCSCXenaTools” R package version 1.4.8.

Kaplan–Meier survival analysis and the Cox Proportional Hazards Model were performed using the “survminer” R package version 0.4.9. The SOX2 expression threshold was calculated using maximally selected rank statistics from the “maxstat” R package version 0.7-25. For the survival curve shown in Figure 6a, the p-values included are the log-rank p-value for the survival probability. We also added the log-rank global p-value for the cox proportional hazard ratio, which is 1.6 with a p of 0.015 for high SOX2 expression. For the survival curve shown in Supplementary Fig6e, the p-value included is the log-rank p-value for the survival probabilities. We also added the log-rank global p-value for the cox proportional hazard ratios, which are 1.5 with a p of 0.032 for high Sox2 expression and 1.6 and p of 0.018 for high TACSTD2 expression.

SOX2 expression in colon and rectal adenocarcinomas (n = 607), and adjacent normal tissue (n = 51) from TCGA, is represented as a density plot in Supplementary Figure S6a. DE analysis was also performed using “DESeq2” R package version 1.38.3. Genes were considered to be differentially expressed if their base mean expression was larger than 2, and their adjusted p-value was lower than 0.05 after correction. The correlation between the expression of SOX2 and TACSTD2 in the colon and rectal adenocarcinoma patients from TCGA was computed using the R package “corrplot” version 0.92 and visualised using the R package “ggpubr” version 0.6.

### Analysis of scRNA-Seq, ATAC-seq and spatial transcriptomics

Raw spatial transcriptome RNA sequencing data was retrieved from GSE225857. Visualisation of gene expression on tissue slices was performed using SpatialFeaturePlot. Data processing and plot generation was performed using the “Seurat” R package version 5.0.3.

Raw single-cell and ATAC-seq data was retrieved form GSE221300. The processing and analysis were performed according to [DOI: 10.1126/sciadv.adf0927]. Briefly, the data was assessed for quality control, normalized, and demultiplexed using the HTODemux function. Principal component analysis was performed, and 10 dimensions with a resolution 0.3 were used to generate the clusters. The data was visualized using Uniform Manifold Approximation and Projection for Dimension Reduction (UMAP) with 10 dimensions. Visualization of the annotation and gene expression was performed using the DimPlot and FeaturePlot functions from Seurat. The processing, quality control, analysis and representation of the scRNA-Seq data was performed using the “Seurat” R package version 4. ATACseq data was loaded in Integrative Genomics Viewer (IGV)^13^ for track visualization and image acquisition.

### CRISPR-Cas9 gene editing

We use nucleofection-based CRISPR-Cas-9 methodology to generate the TP53 KO in the *Sox9^iresCreERT2^* (control) and *Rosa26R-Sox2-IRES-GFP* homozygous (Sox2 HOM) organoid lines. Three separate gRNAs (see Supplementary Table 4 for gRNA sequence) were designed (Synthego ‘Mulit-Guide’ platform), such that their spatial distribution favoured large (>50bp) genomic deletions rather that small indels, resulting in improved knockout efficiency and consistency. sgRNAs were synthesized with improved stability and reduced innate immune responses via 2’-O-methyl analogs on the first and last three bases and 3’ phosphorothioate internucleotide linkages between the first three and last two bases. Ribonucleoprotein (RNP) complexes were formed by diluting 2μl of 100 μM of multi-gRNA (Synthego) in Tris-EDTA (TE) (Synthego) and 1μl of 20μM recombinant Cas-9 (Integrated DNA Technologies (IDT), 1081059) in RNase-free PBS in 12μl Primary Cell P3 Nucleofector solution (Lonza, V4XP- 3032) and incubating at RT for 20 min. To 2.5×10^5^ cells in 5μl Nucleofector solution, 0.8μl of Electroporation Enhancer Solution (IDT, Alt-R Cas9 Electroporation Enhancer, 2 nmol) was added, followed by 15μl of the RNP solution and mixed by pipetting. This was transferred to a well of a 20μl Nucleocuvette Strip (Lonza, V4XP-3032) and transfected using a 4D-Nucleofector Core Unit (Lonza), using the DS-137 program. Cells were allowed to rest for 3 min before being plated in Cultrex BME Type 2 RGF Pathclear and cultured in basal media containing Rho kinase inhibitor Y-27632 (Sigma, Y0503). Media was changed the following day to basal media only. Transfection with 1μl of 20 μM recombinant Cas-9 (Integrated DNA Technologies (IDT), 1081059) in RNase-free PBS in 12μl Primary Cell P3 Nucleofector solution (Lonza, V4XP-3032) was used to generate control cells. To select for TP53 KO clones, control and targeted cells were treated with Nutlin3a (Sigma, #SML0580), a small-molecule inhibitor that stabilizes and activates TP53, leading to cell cycle arrest and apoptosis in cells with functional TP53. Following a 3-day treatment of 10μM Nutlin3a, all the control organoids were dead, whereas the TP53 targeted pool survived in big proportion and were further cultured and expanded in basal conditioned media. Gene knockout was confirmed by WB following UV irradiation to the organoids in order to activate TP53 signalling cascade.

Efficient Sox2 knockout in the AKPT cell line was achieved by electroporation of Cas9-sgRNA ribonucleoprotein (RNP) complexes using a multi-guide approach. For each knockout line, three sgRNAs were designed to simultaneously target an early region of the single coding exon of the *Sox2* gene such that precise repair of the ∼150bp deletion product leads to an out-of-frame edit. Two sgRNA triplets were designed (see Supplementary Table 4 for gRNA sequence). One sgRNA triplet specifically targeted the reported high-mobility group box (HMG) domain of SOX2 (SOX2 KO1). RNPs were formed by combining 0.7μl of each of the three sgRNAs (100 μM in Tris-EDTA, Synthego, custom) in 6μl of SF buffer (Lonza, V4XC-2032) with 1μl recombinant Cas9 2NLS Nuclease (20 μM, Synthego), mixed by vortexing and incubated at room temperature for 30 min. Cultured AKPT cells were detached and dissociated using Accutase (Aldrich, A6964), filtered through a 70 μm strainer and 5×10^5^ cells resuspended in 12μl of SF buffer and 1μL of Electroporation Enhancer solution (IDT, 1076301). 13 μl of the cell suspension was added to the RNP complexes, mixed by pipetting and 20μL transferred to a well of a 20μl Nucleocuvette Strip (Lonza, V4XP-3032). Electroporation was conducted using a 4D-Nucleofector Core Unit (Lonza) and the DS-137 program. Cells were allowed to rest for 3-5 min before being plated and cultured as normal.

### RNA extraction

RNA was extracted according to the manufacturer’s instructions (Qiagen RNeasy, 74106). Harvested cells or organoids were resuspended in RLT buffer (provided with the kit) supplemented with 40mM Dithiothreitol (DTT) to inhibit RNases activity. Before RNA extraction samples were undergone one freeze/thaw cycle to increase the yield of extracted RNA.

### cDNA synthesis and qRT-PCR

500-1000ng of RNA were reverse transcribed using the cDNA Reverse Transcription Kit (Applied Biosystems, #4368813), according to manufacturer’s instructions. RT-qPCR was performed in 384-well plates, in experimental triplicates, in a 12μl reaction mixture containing 6μl of 2x PowerUp™ SYBR® Green Master Mix (Applied Biosystems, A25742), 10μM of each primer (see Supplementary Table 5 for primer details) and 25-50ng of cDNA. The reaction mixture without cDNA template was used as a negative control for each reaction plate. After 40 cycles of amplification, samples were normalised to the housekeeping gene *Ppib*, where data was expressed as mean ± s.e.m.

### Statistical Analysis

Results are expressed as mean ± standard error of the mean (∗P < .05, ∗∗P < .01, ∗∗∗P < .001, ∗∗∗P < .0001). Statistical significance of mean values was assessed using unpaired Student t-test or analysis of variance, 1- or 2-way, followed by Tukey’s or Sidak’s Multiple Comparison Post-test respectively. The corresponding number of N and experiments are indicated in the figure legends. Statistics were performed using GraphPad Prism 7 software (La Jolla, CA).

---

**Fig. S1.**
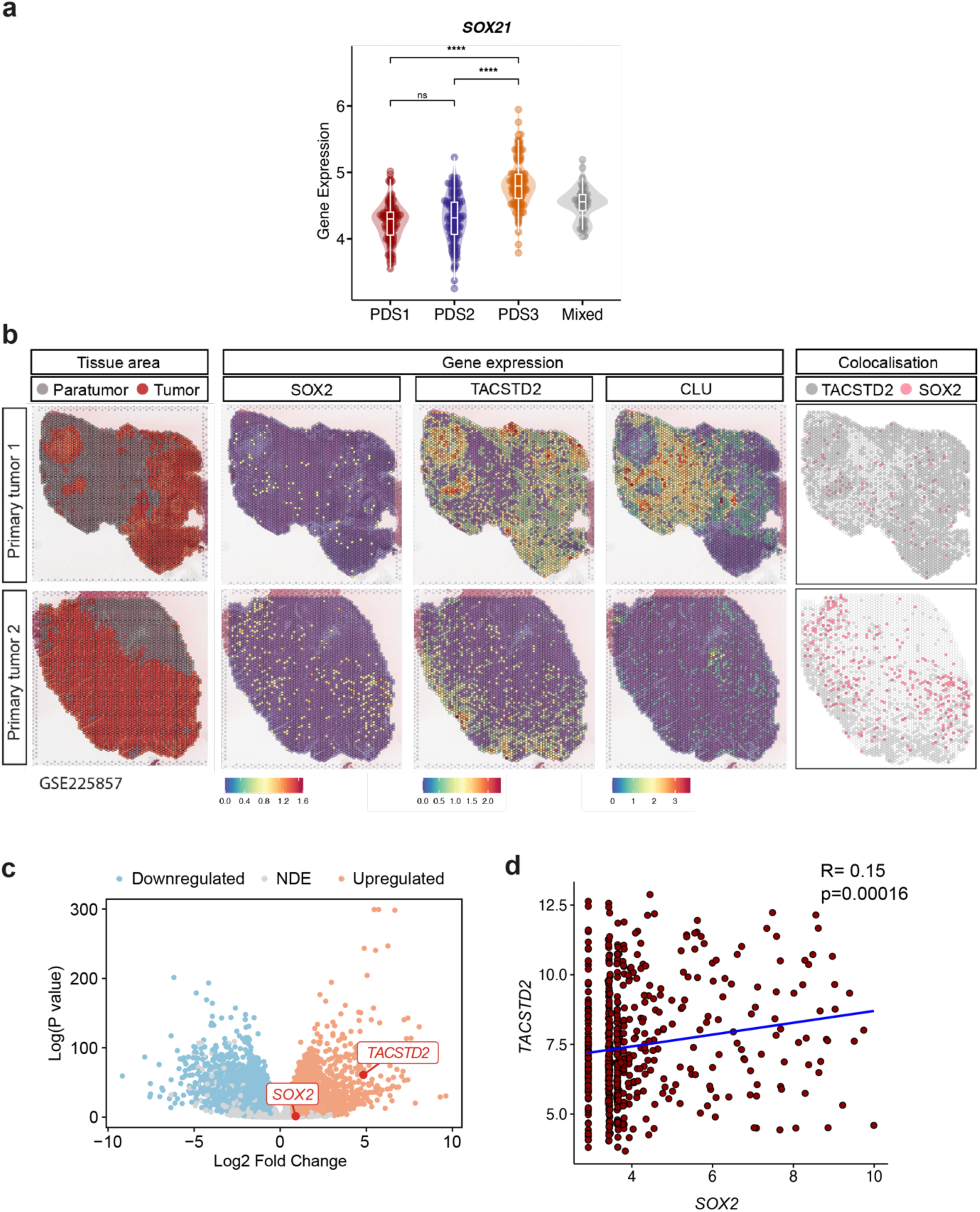
SOX2 expression correlates with reprogramming in human CRC. **a**, Violin plot displaying *SOX21* expression across PDS in the FOCUS cohort (GSE156915; n=360). **b**, Spatial transcriptome RNA sequencing data visualization of SOX2, TACSTD2 and CLU in two primary colorectal tumours. Tumour and paratumour areas were segmented by clustering and annotated using expression markers (GSE225857). **c**, Volcano plot showing significantly upregulated (orange) or downregulated (blue) differentially expressed genes in normal vs tumour COAD and READ RNAseq samples from TCGA. **d,** Correlation plot between *TACSTD2* and *SOX2* in COAD and READ samples from TCGA.

**Fig. S2.**
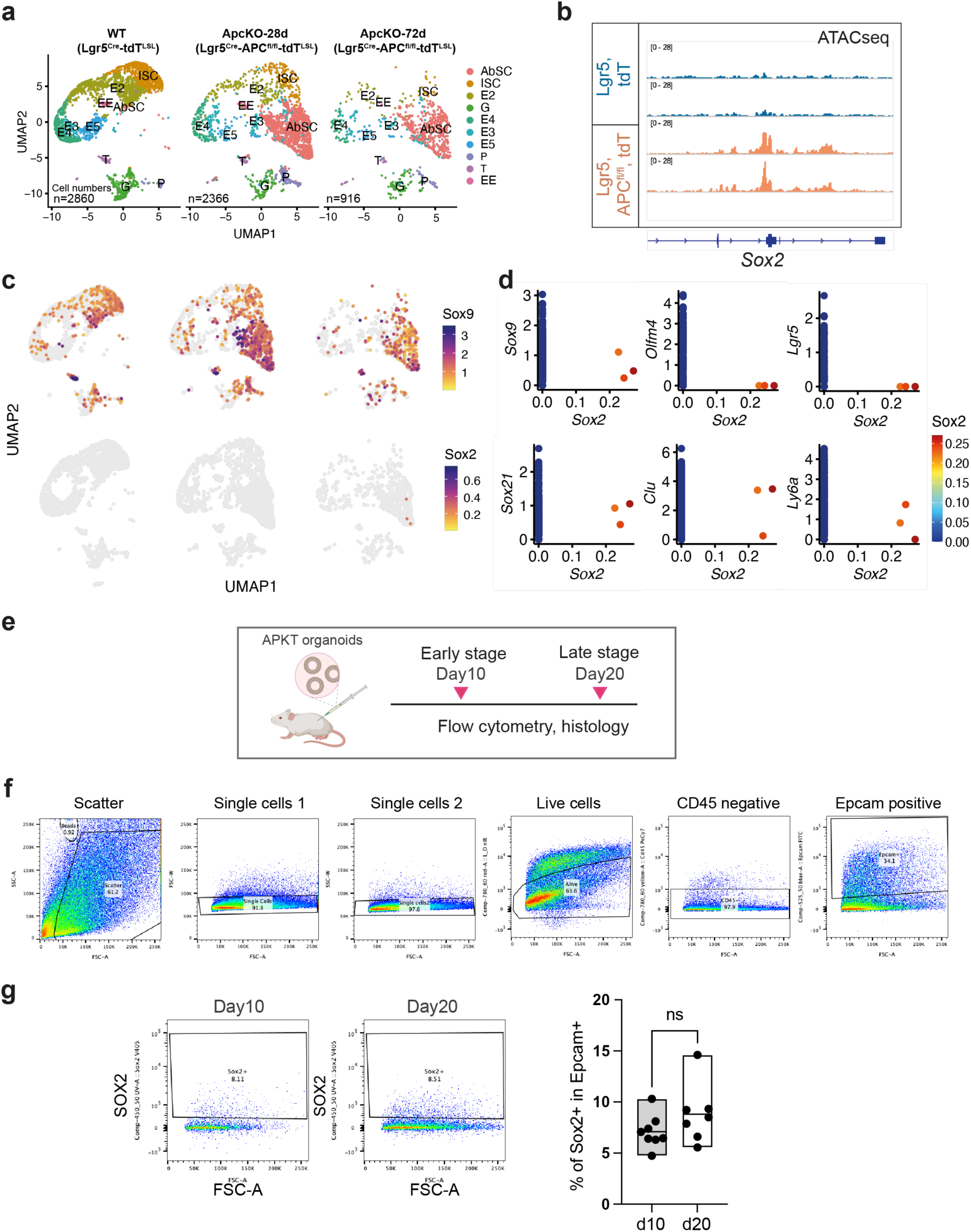
SOX2+ cells are detected in both early adenomas and advanced CRC. **a,** scRNAseq data re-analysis from GSE221300 (Bala et al, 2023) showing the appearance of the aberrant stem cell population (AbSC) upon APC loss following tamoxifen induction at day 28 and 72. **b,** ATACseq data from GSE221300 (Bala et al, 2023) showing increased chromatin accessibility in *Sox2* locus following APC loss. **c,** UMAPS from the scRNAseq re-analysis (GSE221300) showing *Sox9* and *Sox2* expression in WT, ApcKO 28 days and ApcKO 72 days. **d,** *Sox9*, *Olfm4*, *Lgr5*, *Sox21*, *Clu* and *Ly6a* expression of Sox2+ cells detected in ApcKO-72days sample shown in (c). **e,** Scheme showing the experimental design of APKT organoids xenograft mouse model for flowcytometry and histology analyses. **f,** Gating strategy of APKT dissociated tumours for flowcytometry. **g,** Representative flowcytometry plots showing Sox2+ positive population in dissociated single cells from APKT tumours collected at day 10 and day 20. Cells were gated following sequential selection (as indicated in f) and Epcam+ cells were plotted by Sox2 exclusion vs SSC. On the right, quantification of percentage of Sox2+ cells present in the Epcam+ population at day 10 (n=8) and day 20 (n=7). Data represent mean ± s.e.m. ns, not significant, unpaired two-sided t-test.

**Fig. S3.**
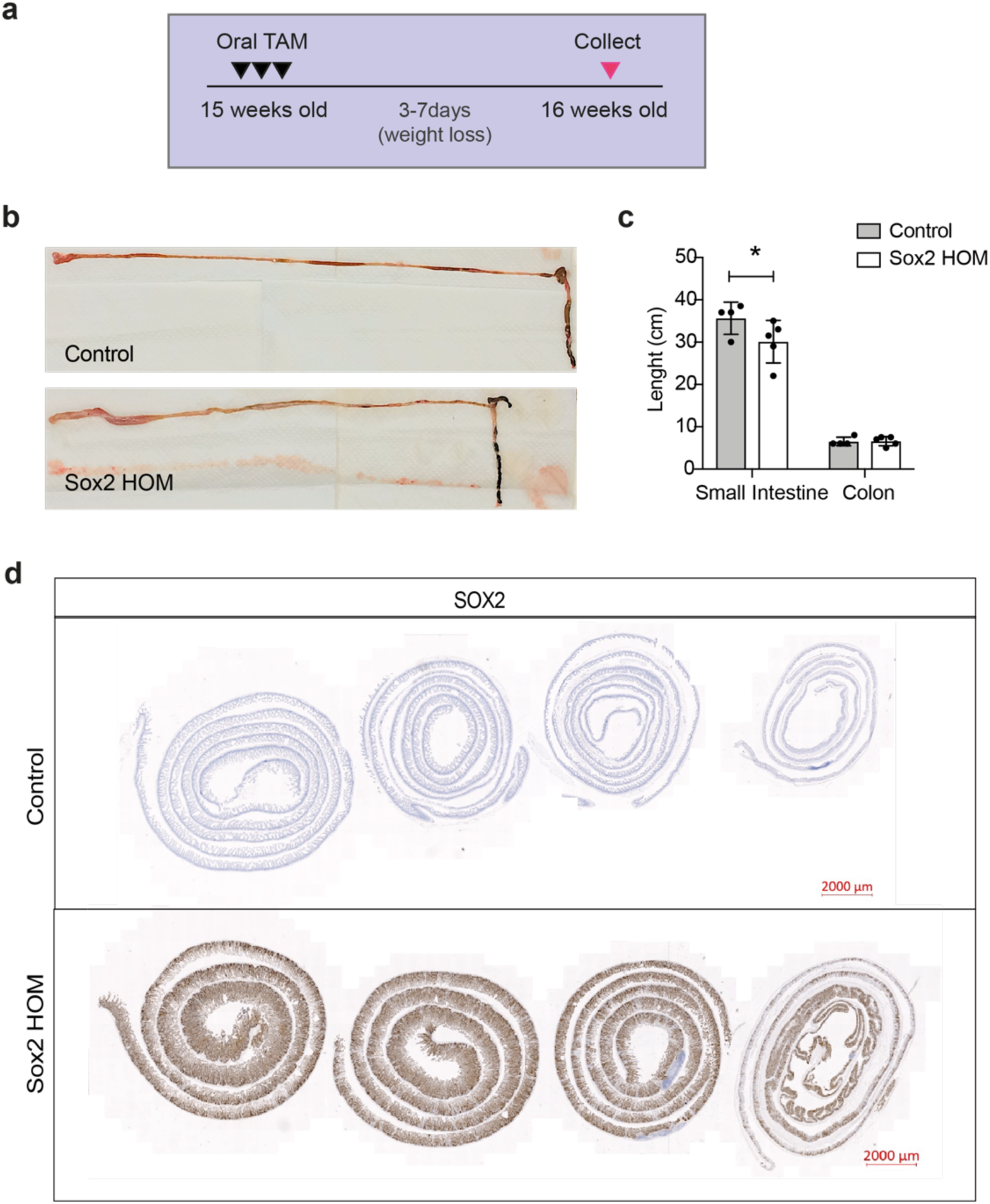
Homozygous SOX2 expression induces acute intestinal hyperplasia. **a,** Scheme of tamoxifen administration to control and Sox2 HOM mice. **b,** Macroscopic appearance of the gut isolated from control and Sox2 HOM mice 1 week after tamoxifen administration. **c,** Length measurements of the small intestine and colon of these animals. Data represent mean ± s.e.m. *P<0.05, **P<0.01, ***P<0.001, 2-way ANOVA and Šídák’s multiple comparisons test. **d,** SOX2 IHC in control and Sox2 HOM mice 1 week after TM induction. Scale bar, 2000μm.

**Fig. S4.**
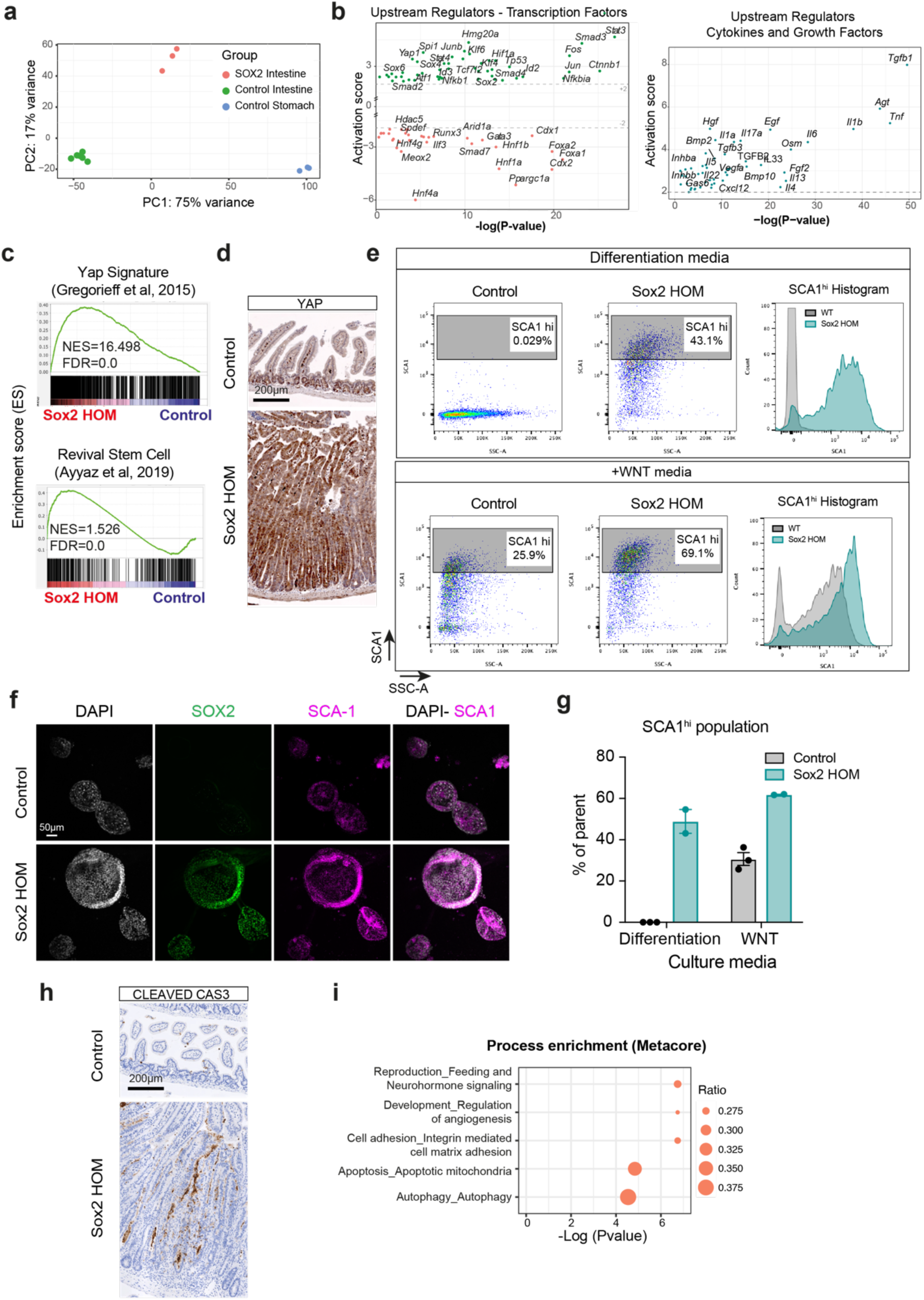
Homozygous SOX2 expression induces fetal reprogramming. **a,** Principal component analysis (PCA) plot of transcriptomic data from control intestine (n=6), control stomach (n=3) and Sox2 HOM intestine (n=3). **b,** Plot showing statistically significant activated or inhibited (activation z-score) predicted upstream regulators of the enriched pathways from Sox2 HOM intestine (n=3) vs control (n=6) intestines. Transcription factors (above) and cytokines and growth factors (below). **c,** GSEA of revival stem cell and Yap signatures. **d,** IHC of YAP in control and Sox2 HOM intestines 7 days following TM administration. Scale bar, 200μm. **e,** Representative flowcytometry plots and histograms of dissociated single cells from control and Sox2 HOM organoids cultured in WNT or differentiation media. Cells were gated following sequential selection by SCA1 exclusion vs SSC. **f,** Confocal immunofluorescence for SOX2 and SCA1 in control and Sox2 HOM organoids cultured in WNT media. Scale bar, 50μm. **g,** Percentage of control or Sox2 HOM cells expressing high levels of SCA1 in WNT and differentiation media culture conditions. Data represent mean ± s.e.m. of biological replicates. **h,** IHC of Cleaved Caspase3 in control and Sox2 HOM intestines 7 days following TM administration. Scale bar, 200μm. **i,** Metacore top 5 significant enriched processes in Sox2 HOM vs control intestines. Scale bar, 200μm.

**Fig. S5.**
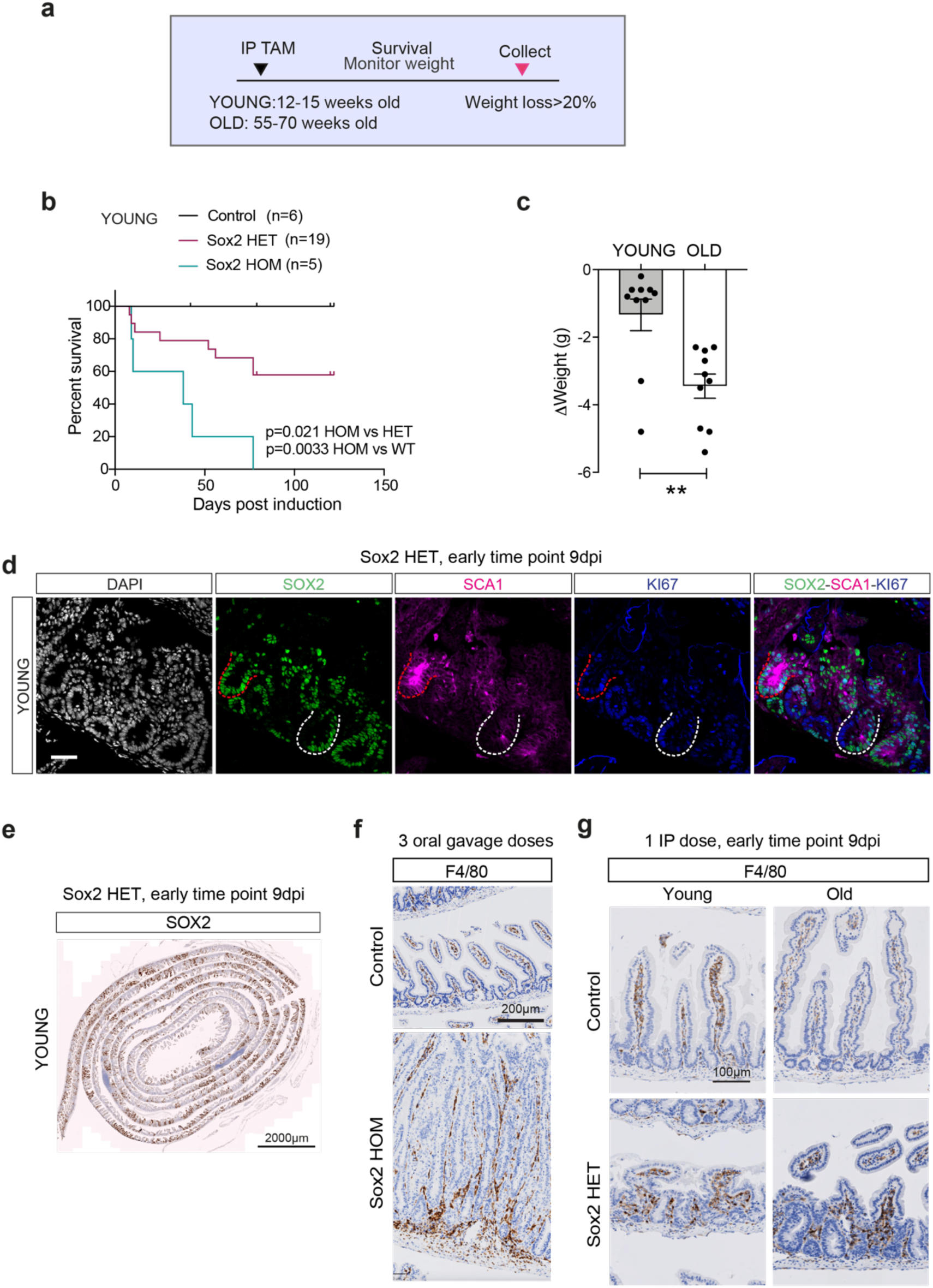
SOX2-induced reprogramming triggers immune surveillance. **a,** Scheme of tamoxifen administration and study design for the survival cohort of young and old control, Sox2 HET and Sox2 HOM mice. **b,** Kaplan-Meier survival analysis of Control, Sox2 HET and Sox2 HOM young animals following one i.p. dose of TM. p-values were determined using the Mantel-Cox test (P<0.05, **P<0.01, ***P<0.001). **c,** Bar chart representing weight increase at the time of death in young and old Sox2 HET mice after tamoxifen treatment. Data represent mean ± s.e.m. **P<0.01, unpaired two-sided t-test test. **d,**Triple immunofluorescence for SOX2, SCA1 and KI67 in young Sox2 HET mice intestinal sections at 9dpi. Scale bar, 200μm. **e,** IHC of SOX2 in young Sox2 HET mice intestinal sections at 9dpi. Scale bar, 2000μm. **f,** IHC of F4/80 (macrophages) in control and Sox2 HOM intestinal sections following TM administration as indicated. Scale bar, 200μm. **g,** IHC of F4/80 (macrophages) in control and Sox2 HOM intestinal sections of young and old mice following TM administration as indicated. Scale bar, 100μm.

**Fig. S6.**
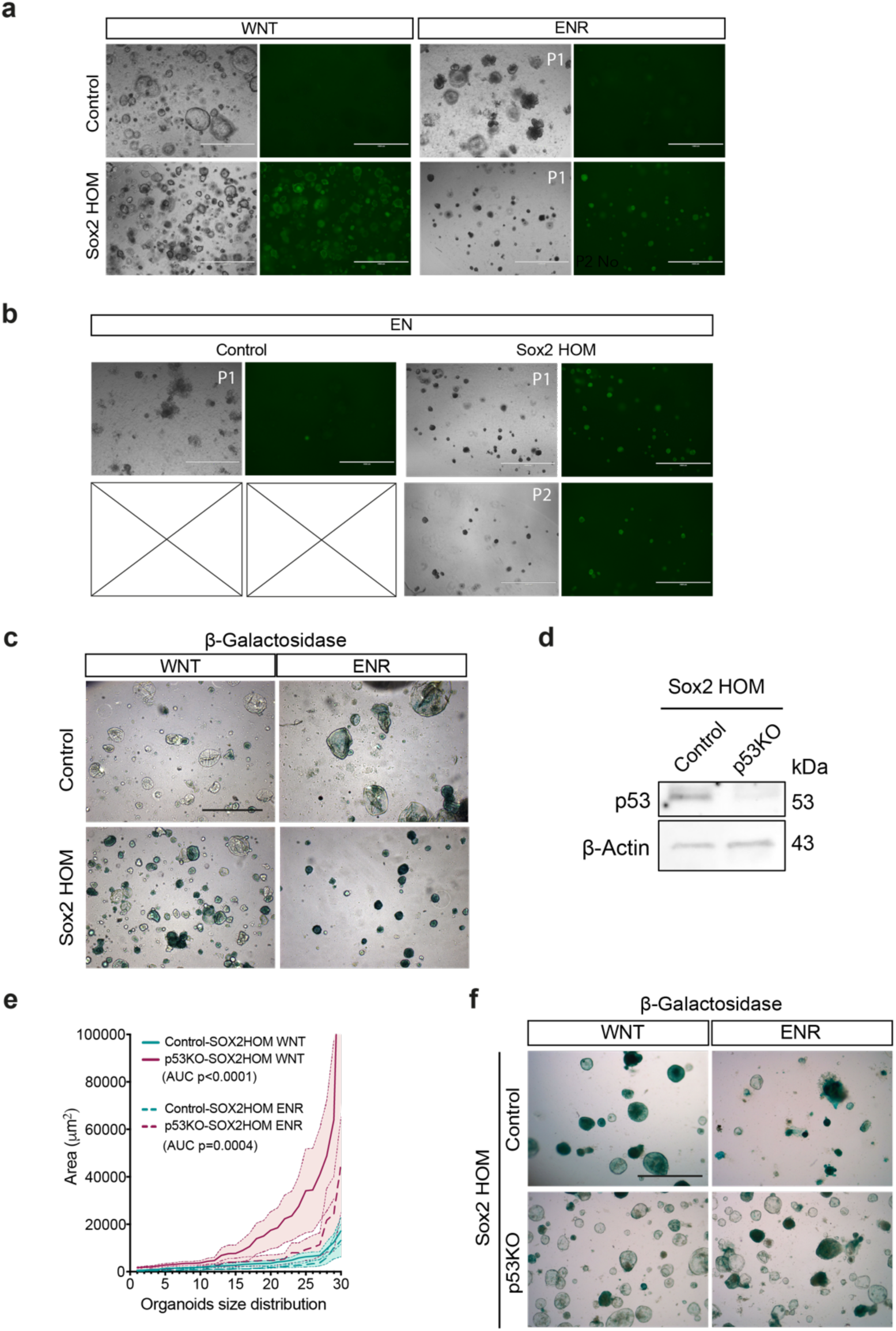
Loss of p53 reverses SOX2-induced senescence. **a,** Representative images of organoids derived from control and Sox2 HOM intestines cultured in WNT media (left panel) and transferred to ENR media (picture taken after one passage, P1). Scale bar, 1000μm. **b,** R-spondin withdrawal challenge (EN media) one passage after transferring them from WNT media culture condition (passage 1, P1) and maintained for a second passage on EN culture conditions (passage 2, P2). Cross indicates the conditions could not be maintained. Scale bar, 1000μm. **c,** β-Galactosidase staining in control and Sox2 HOM organoids cultured in WNT or ENR media conditions. Scale bar, 1000μm. **d,** Western blot analysis of p53 protein levels in control (Cas9)-Sox2 HOM and p53KO-Sox2 HOM intestinal organoids. **e,** Organoid size distribution by area measurement in control (Cas9)-Sox2 HOM (green) and p53KO-Sox2 HOM (red) intestinal organoids cultured in WNT (thick line) or ENR media (dotted line). Average of 3 independent experiments, ***P<0.001, ****P<0.0001, AUC. **f,** β-Galactosidase staining in control Sox2 HOM (Cas9) and Sox2 HOM p53KO intestinal organoids cultured in WNT or ENR media conditions. Scale bar, 1000μm.

**Fig. S7.**
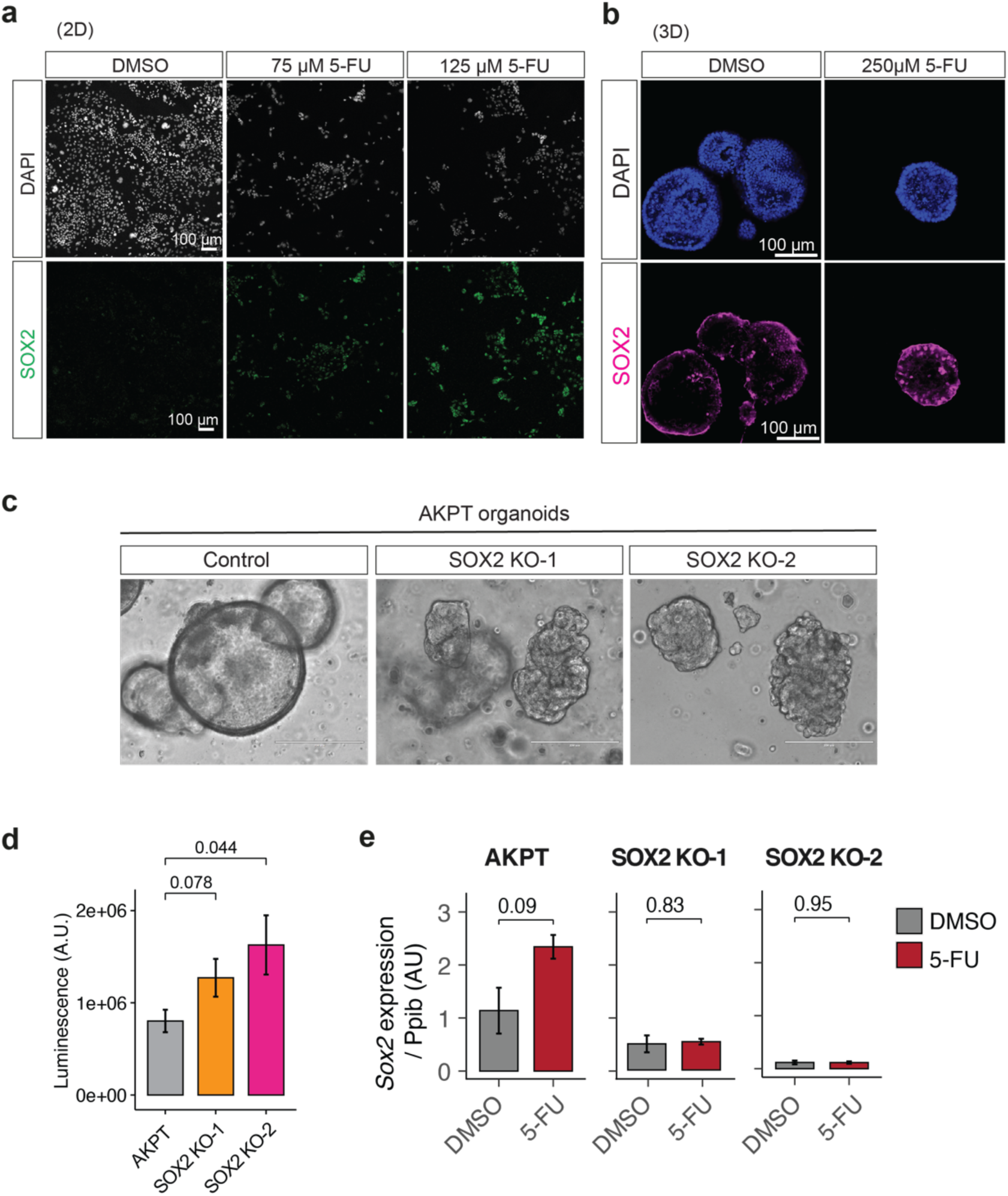
Sox2 deletion suppresses fetal reprogramming and cystic morphology in the AKPT CRC organoids. **a,** Representative immunofluorescence of Sox2 and DAPI in the VAKPT cell line. The cells were grown in glass slides, fixed, stained and imaged using confocal microscopy. **b**, Representative immunofluorescence of SOX2 in AKPT organoids treated with 250 µM 5-FU and DMSO as a control. The organoids were fixed, permeabilized and stained with anti-Sox2, and DAPI. Scale bar: 100 µM. **c**, Representative phase contrast images of WT and the two Sox2 KO models of AKPT organoids. Scale bar: 200 µm. **d**, Proliferation of WT and two Sox2 KO models of AKPT cell lines. The cells were seeded and left to proliferate for 72 hrs. The number of viable cells is proportional to the luminescent signal. The Welch Two Sample t-test was performed to test to compare the WT to the Sox2 KO cells. **e**, qPCR quantification of Sox2 in WT and two Sox2 KO AKPT organoids treated with 5-FU (250 µM), or DMSO for 72 hrs. Expression values are normalized to Ppib levels and represented as Log2 Fold Change. The experiment was performed in three biological replicates, and each sample was measured in three technical replicates for the qPCR quantification. Welch Two Sample t-test was used to test the differences between the DMSO and the 5-FU treatment for each of the organoid lines.

